# Activation of ATF3 via the Integrated Stress Response Pathway Regulates Innate Immune Response to Restrict Zika Virus

**DOI:** 10.1101/2023.07.26.550716

**Authors:** Pheonah Badu, Gabriele Baniulyte, Morgan A. Sammons, Cara T. Pager

## Abstract

Zika virus (ZIKV) is a re-emerging mosquito-borne flavivirus that can have devastating health consequences. The developmental and neurological effects from a ZIKV infection arise in part from the virus triggering cellular stress pathways and perturbing transcriptional programs. To date, the underlying mechanisms of transcriptional control directing viral restriction and virus-host interaction are understudied. Activating Transcription Factor 3 (ATF3) is a stress-induced transcriptional effector that modulates the expression of genes involved in a myriad of cellular processes, including inflammation and antiviral responses, to restore cellular homeostasis. While ATF3 is known to be upregulated during ZIKV infection, the mode by which ATF3 is activated and the specific role of ATF3 during ZIKV infection is unknown. In this study, we show via inhibitor and RNA interference approaches that ZIKV infection initiates the integrated stress response pathway to activate ATF4 which in turn induces ATF3 expression. Additionally, by using CRISPR-Cas9 system to delete ATF3, we found that ATF3 acts to limit ZIKV gene expression in A549 cells. We also determined that ATF3 enhances the expression of antiviral genes such as STAT1 and other components in the innate immunity pathway to induce an ATF3-dependent anti-ZIKV response. Our study reveals crosstalk between the integrated stress response and innate immune response pathways and highlights an important role for ATF3 in establishing an antiviral effect during ZIKV infection.

**Importance:** ZIKV is a re-emerging mosquito-borne flavivirus that co-opts cellular mechanisms to support viral processes which can reprogram the host transcriptional profile. Such viral-directed transcriptional changes and the pro- or anti-viral outcomes remain understudied. We previously showed that ATF3, a stress-induced transcription factor, is significantly upregulated in ZIKV infected mammalian cells, along with other cellular and immune response genes. We now define the intracellular pathway responsible for ATF3 activation and elucidate the impact of ATF3 expression on ZIKV infection. We show that during ZIKV infection the integrated stress response pathway stimulates ATF3 which enhances the innate immune response to antagonize ZIKV infection. This study establishes a link between viral-induced stress response and transcriptional regulation of host defense pathways and thus expands our knowledge on virus-mediated transcriptional mechanisms and transcriptional control of interferon stimulated genes during ZIKV infection.

## Introduction

Zika virus (ZIKV) is a flavivirus that is spread mainly by *Aedes* mosquitoes (1) and causes self-limiting infections characterized by mild symptoms such as fever, headache, and joint pain (2). The re-emergence of ZIKV from 2007 to 2016 produced large outbreaks on the Yap Island, French Polynesia, and the American region (3). These outbreaks implicated the virus in intrauterine-linked complications termed congenital Zika syndrome which includes microcephaly, congenital malformations, and fetal demise (4–6). Additionally, the recent surges in infection in adults also revealed an association with Guillain-Barré syndrome, a neurological disease which results in paralysis (7–10). Combined these damaging effects make re-emerging ZIKV a significant public health challenge, which is worsened by climate-induced vector expansion, mosquito, sexual and intrauterine transmission routes (11–15) and the absence of antiviral drugs and vaccines. Improving our understanding of the core mechanisms of viral processes, virus-host interactions, and viral restriction may provide valuable clues to help offset this re-emerging public health challenge.

ZIKV has a single-stranded positive-sense RNA genome, approximately 11,000 nucleotides in length, that is translated into a single polyprotein upon viral entry into a host cell. Viral translation occurs on the endoplasmic reticulum (ER) membrane and is followed by proteolytic cleavage of the polyprotein. This process produces structural proteins (capsid [C], precursor membrane [prM], and envelope [E]) involved in formation of virions and non-structural proteins required for protein processing (NS2B and NS3), viral replication (NS1, NS2A, NS3. NS4A, NS4B, and NS5, the RNA dependent RNA polymerase [RdRp]), and immune evasion (NS1, NS5) (16, 17). After these viral proteins are made, the viral genome is replicated on the ER membrane. This process triggers extensive remodeling of the ER membrane as host proteins together with viral nonstructural (NS) proteins assemble to form replication complexes (18, 19). Newly replicated genomes subsequently associate with structural proteins to form the nascent virion on the ER membrane at sites juxtaposed to the replication complex (18, 19). As a result of the immense structural changes to the ER membrane and the accumulation of misfolded proteins in the ER, cellular homeostasis is disrupted. In response, the cell activates two distinct but overlapping signaling networks namely the Unfolded Protein Response (UPR) and the Integrated Stress Response (ISR) (20–23).

The ISR is a network of signaling pathways in eukaryotic cells stimulated by external and internal stressors including viral infection, nutrient deprivation, and ER stress (24). These stressors activate a four-member family of eIF2⍺ kinases, PERK (Protein Kinase R-like ER kinase), PKR (Protein Kinase R; a double-stranded RNA-dependent protein kinase), GCN2 (general control non-derepressible-2) and HRI (heme-regulated eIF2⍺ kinase) (25). All four kinases share sequence similarity in their catalytic domains but have different regulatory domains. Therefore, each kinase responds to a distinct stress, but all target the translation initiation factor eIF2 and phosphorylate the serine 51 residue of the alpha subunit (26). This phosphorylation event inhibits the guanine nucleotide exchange factor for the eIF2 complex, eIF2B, and prevents the assembly of translation pre-initiation complexes (26). Ultimately, eIF2⍺ phosphorylation represses global cap-dependent translation but promotes the preferential translation of select mRNAs that play key roles in resolving the stress (27).

Activating transcription factor 4 (ATF4) is one of the best studied effector proteins of the ISR (20, 23). This transcription factor acts as a master regulator of stress and is selectively translated through a mechanism involving the delayed translational reinitiation on an upstream open reading frame upon eIF2⍺ phosphorylation (27, 28). When induced, ATF4 controls the transcriptional programs of a cohort of genes involved in cell survival or cell death. The overall outcome of ATF4 expression is context specific and is influenced by the cell type, type of stressor, and the duration of stress (29, 30). One target of ATF4 is Activating Transcription Factor 3 (ATF3), another stress response gene activated during stressful conditions. Depending on the cellular environment or nature of the stress, ATF3 can be activated by other effectors beside ATF4 (31–33). Like ATF4, ATF3 belongs to the ATF/CREB family of transcription factors and can function as either a transcriptional activator or repressor (31–33). ATF3 has a DNA binding domain as well as a basic leucine zipper (bZip) region that is important for dimer formation (34). When promoting transcription of target genes, ATF3 heterodimerizes with other bZip proteins like c-JUN, while in a repressive role, ATF3 forms homodimers or stabilizes inhibitory co-factors at promoter sites (34, 35). Generally, ATF3 modulates various cellular processes like autophagy, innate immune and inflammatory responses, DNA damage response, and cell cycle progression (31–33). During viral infection, activation of ATF3 produces paradoxical outcomes (36–38). Notably during infection with the mosquito-borne flavivirus Japanese encephalitis virus (JEV), ATF3 was shown to repress the expression of select interferon stimulated and autophagy genes which enhanced viral protein and RNA levels (37).

Our recent global transcriptomic analysis of human neuronal SH-SY5Y cells infected two different isolates of ZIKV, Uganda (MR799) and Puerto Rico (PRVABC59), and DENV serotype 2 revealed an upregulation of immune response genes in both ZIKV strains but not in DENV (39). Additionally, genes involved in cellular responses were significantly upregulated particularly in PRVABC59 infected cells, including genes associated with both the UPR and ISR pathways (e.g., *ATF4*, *ATF3,* and *CHOP/DDIT3*) (39). Elevated *ATF4* expression indicated that the ISR pathway was activated during ZIKV PRVABC59 infection, which in turn stimulated *ATF3* expression and downstream targets like *CHOP* for stress management. However, the functional significance of ATF3 in ZIKV infection and if this stress-induced transcription factor exhibited pro- or anti-viral functions, had not been determined.

In this study, we used ISR-specific inhibitors and RNAi approaches to show that during ZIKV infection the ISR pathway stimulated ATF4 expression which directly activated ATF3. We further demonstrated that in the absence of ATF3, the levels of ZIKV protein, RNA, and virions increased, indicating that ATF3 functioned to restrict viral infection. Finally, we determined that knockout of ATF3 altered the expression of anti-viral innate immune genes during ZIKV infection. Our data reveal the effects of ATF3 regulation within the cell and highlight that ATF3-driven regulation of innate immunity pathways impedes ZIKV infection.

## Results

### ZIKV induces strong ATF3 expression 24-hours post infection

In a previous gene expression study, we observed that ZIKV PRVABC59 (ZIKV^PR^) infection in a neuronal cell line (SH-SY5Y) stimulated immune and stress response genes such as *ATF3* and *CHOP* (39). Moreover, in a reanalysis of RNA-seq data collected from peripheral blood mononuclear cells from patients in early- and late-acute and convalescent stages of ZIKV infection, we determined that *ATF3* levels were increased (40). To determine when ATF3 was stimulated during ZIKV infection, we infected cells with ZIKV and examined viral and cellular proteins and RNA levels at different timepoints following infection. In this research we used the human A549 lung adenocarcinoma cell line as these cells support robust ZIKV infection (41–44), can induce an immune response upon viral infection (42, 44), and are a tractable cell culture system to investigate foundational molecular mechanisms and cellular pathways influencing ZIKV infections (40). The highest level of the ZIKV nonstructural protein NS1 was observed at 24 hours post-infection and correlated with peak ATF3 protein expression (Fig 1A). ATF4 protein expression increased from 12- to 24-hours following infection and remained steady until 48 hours post-infection (Fig 1A). Consistent with this trend, viral, *ATF4*, *ATF3,* and *CHOP* mRNA significantly increased at 24 hours post-infection (Fig 1B-E). Since high viral protein and RNA production occurred at 24 hours post-infection, we reasoned that translation and replication peaked 24 hours after ZIKV infection and declined by 48 hours as virion packaging occurred. As predicted, a high titer of virions was released 48 hours after infection (Fig 1F). We similarly examined ATF3 expression following infection with MR766, the original ZIKV strain isolated in Uganda in 1947 (1, 45). ZIKV MR766 also induced ATF3 mRNA and protein expression, albeit at 48 hours post-infection compared to 24 hours for ZIKV PRVABC59 (data not shown). Together, these data indicated that peak viral protein and RNA expression coincided with ATF3 RNA and protein expression. Moreover, the induction of ATF3 expression during ZIKV infection is consistent with increased ATF3 levels in two biologically relevant systems to ZIKV infection namely SH-SY5Y neuronal cells and peripheral blood mononuclear cells (PBMCs) isolated from ZIKV-infected patients (39, 40).

**FIG 1.**
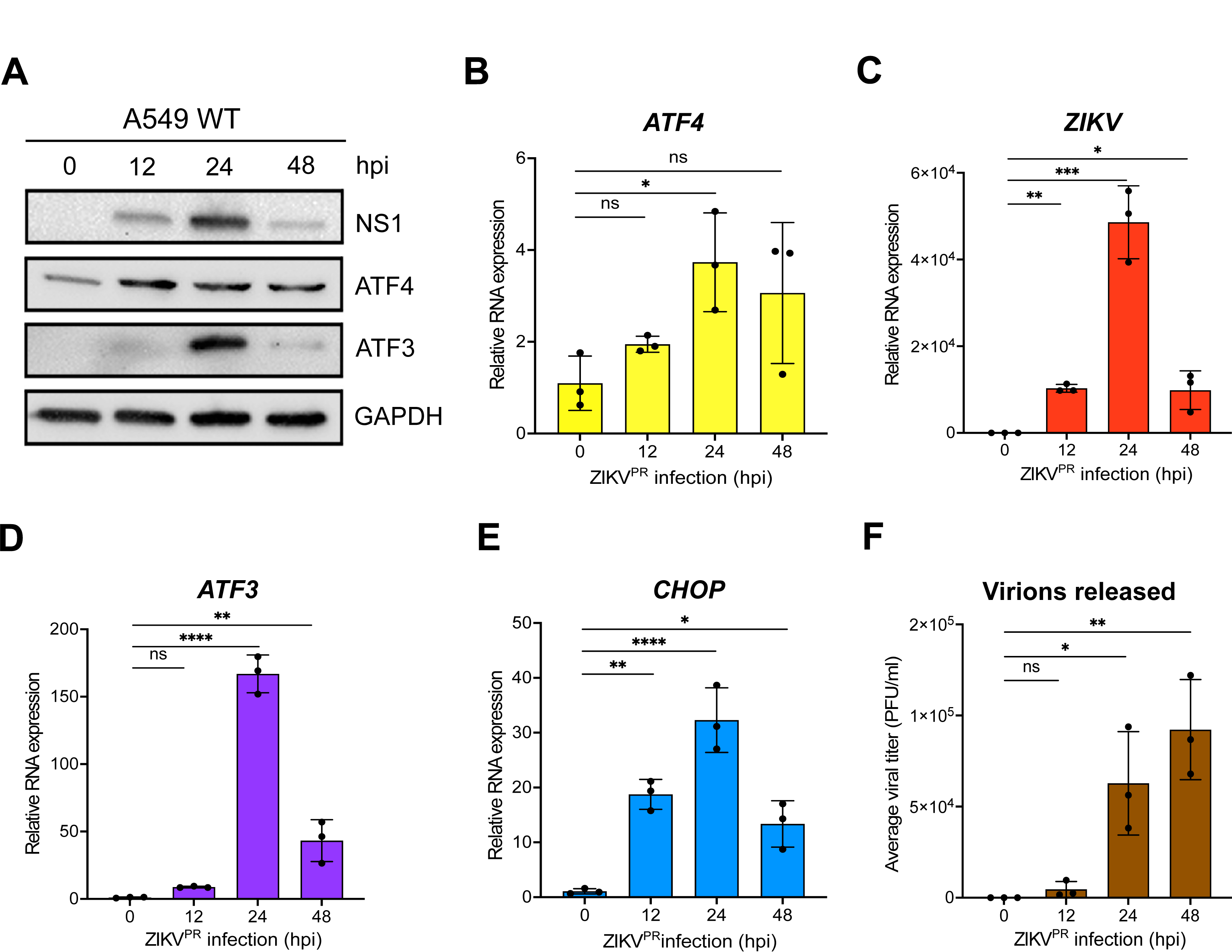
ZIKV significantly induces ATF3 expression 24-hours after infection. (**A**) A549 WT cells were infected with ZIKV^PR^ at moi of 10 PFU/cell for 0-, 12-, 24- or 48-hours post-infection (hpi). Cellular (ATF4 and ATF3) and viral (NS1) proteins were assayed by western blot with GAPDH as the loading control. Western blot is representative of at least three independent experiments. (**B-E**) Total RNA was extracted at the indicated timepoints and used as template for RT-qPCR to measure the expression of *ATF4*, ZIKV, *ATF3* and *CHOP* mRNAs. The relative mRNA expression was determined by the 2^−ΔΔCt^ method using mock-infected cells as reference and the genes were normalized to *ACTB*. RT-qPCR data are means ± SD of three technical replicates and determined from three independent experiments. (**F**) Viral titers in the cell culture media collected at the different infection time points were measured by plaque assay. PFU, plaque forming units. The data represent the means ± SD of two technical replicates from three independent experiments. Statistical significance was determined by Student T-test. *p<0.01, **p<0.001, ***p<0.0005, ****p<0.0001, ns-not significant.

### ATF3 restricts ZIKV gene expression

To determine the functional importance of ATF3 during ZIKV infection, we generated an ATF3 knock-out (KO) A549 cell line using CRISPR-Cas9 gene editing with a guide RNA targeting exon 2. We validated ATF3 KO by sequence analysis (data not shown) and by comparing ATF3 expression in WT and KO cell lines treated with DMSO or tunicamycin (data not shown). Tunicamycin inhibits the first step of protein N-linked glycosylation to affect the folding of glycosylated proteins in the ER (46, 47). The accumulation of these misfolded proteins in the ER lumen induces ER stress, activation of PERK, a UPR and ISR sensor, which phosphorylates eIF2⍺ and enhances translation of ATF4 to induce ATF3 expression. Indeed, in WT A549 cells ATF3 expression was induced by tunicamycin treatment, but ATF3 protein was absent in the KO cells (data not shown). Notably, RT-qPCR analysis showed that ATF3 mRNA was upregulated in the KO cells (data not shown). Because the gRNA used to generate the KO cells targets a region within exon 2 which contains the translational start codon, transcription of *ATF3* was not affected by genome editing, whereas expression of the ATF3 protein was strongly inhibited (data not shown). Hence, when ATF4, the upstream effector of ATF3, was induced upon stress, the effector activated the transcription of ATF3, but downstream translation was impeded.

Next, WT and ATF3 KO cells were mock-infected or infected with ZIKV at two different moi (1 and 10 PFU/cell). Cells were harvested at 24 hours post-infection, and virus and ATF3 expression examined by western blotting and RT-qPCR. Our data showed that ZIKV infection induced ATF3 protein expression in WT cells but not in ATF3 KO cells (Fig 2A). Interestingly, we found that in ATF3 deficient cells the levels of the ZIKV NS1 protein were notably increased compared to NS1 levels in WT cells (Fig 2A). Consistent with the increase in ZIKV protein, viral RNA was significantly upregulated in ATF3 deficient cells compared to WT cells (Fig 2B). Consistent with the tunicamycin treated cells (data not shown), ATF3 protein and RNA expression was induced by infection in WT cells and absent in the ATF3 KO cells (Fig 2A and Fig 2C). We additionally performed plaque assays to quantify virion titer produced in WT and ATF3 KO cells and determined that a greater number of infectious particles were produced in the absence of ATF3 (Fig 2D). To validate these data, we also examined ZIKV gene expression in WT HCT-116 colorectal cells, which have high ATF3 expression profile (48), and ATF3 KO HCT-116 cells, which were generated by an alternative gene editing approach based on adeno-associated virus mediated homologous recombination (49). We observed a similar increase in the level of ZIKV protein, RNA, and viral titers in infected ATF3 KO HCT116 cells (Fig 2E-G). Overall, these results indicate that ATF3 expression suppressed ZIKV gene expression, and this effect was not cell type specific.

**Fig 2.**
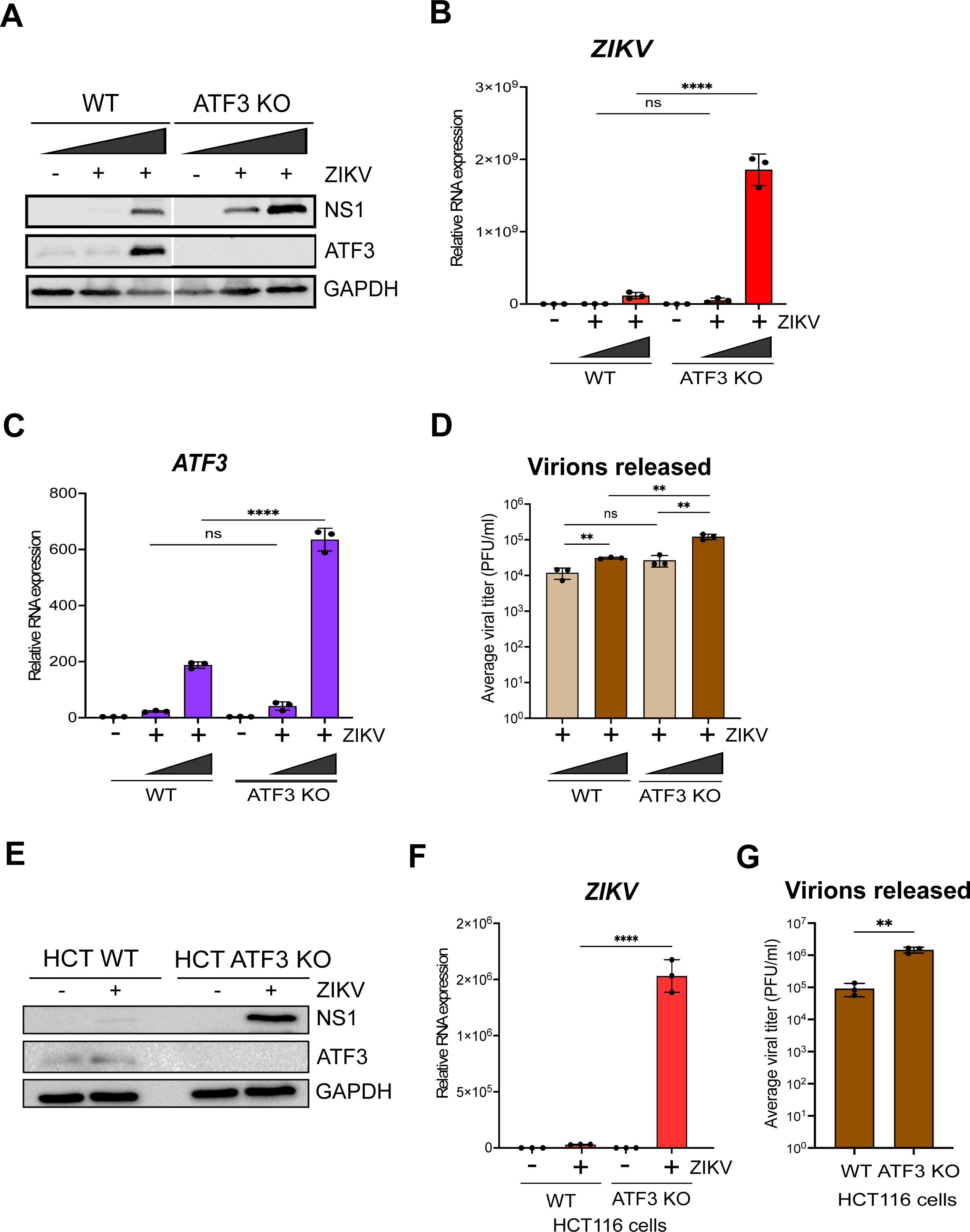
ATF3 restricts ZIKV gene expression. The effect of ATF3 expression on ZIKV infection was examined by infecting A549 WT and ATF3 KO cells with or without ZIKV^PR^ (moi of 1 and 10 PFU/cell) for 24 hours. (**A**) ZIKV NS1 and ATF3 proteins were analyzed by western blot in which GAPDH was used as the loading control. The western blot shown is a representative of three independent experiments. Total RNA from infected cells were analyzed by RT-qPCR using primers specific to (**B**) ZIKV and (**C**) *ATF3.* The RNA expression was normalized to *ACTB* and the relative transcript expression was calculated by the 2^−ΔΔCt^ method using mock-infected cells as reference for the different cell lines. The data shown are means ± SD for three technical replicates and are from three independent experiments. (**D**) Virions released during infection in WT and ATF3 KO cells were quantified as the average viral titer (PFU/ml) using the plaque assay method. The mean PFU/ml ± SD was derived from two technical assays of three independent experiments. (**E-F**) To validate the role of ATF3 in ZIKV infection, HCT-116 WT and ATF3 KO cells were infected with ZIKV^PR^ (moi=10 PFU/cell) for 48 hours. (**E**) ATF3 and viral NS1 proteins were analyzed by western blot with GAPDH as the loading control. (**F**) ZIKV RNA expression normalized to *ACTB* was determined by RT-qPCR (2^−ΔΔCt^ method). Data are from three independent experiments and three technical replicates within each experiment and are shown as mean ± SD of. (**G**) Viral titers in HCT-116 cell culture media were measured by plaque assay. Statistical significance was determined by Student T-test. **p<0.05, ****p<0.0001, ns-not significant

### ATF3 is activated through the ISR pathway during ZIKV infection

A number of effector proteins (e.g., ATF4, p53, NF-kB, and JNK), associated with different signaling pathways, are known to induce ATF3 expression (31–33). Given that ZIKV induces changes in ER membrane morphology (19), activates ER stress sensors (IRE-1, ATF6, and PERK) (41, 50, 51), and the presence of double-stranded viral RNA intermediates activate PKR (41, 52, 53), we reasoned that increased ATF3 expression was initiated through the ISR pathway (Fig 3A). Specifically, activation of the ISR kinases during ZIKV infection would lead to a shutdown of cap-dependent translation, increase translation of ATF4, and subsequent activation of ATF3 (Fig 3A). To investigate if the ISR pathway was responsible for ATF3 activation during ZIKV infection, we inhibited the ISR pathway in mock- and ZIKV-infected cells using a general ISR inhibitor (ISRIB). ISRIB acts on eIF2B, a guanine nucleotide exchange factor involved in translation and renders the cells resistant to the effects of eIF2⍺ phosphorylation (54–56). ISRIB or DMSO (vehicle control) were added to cells 1-hour after the initial virus infection and maintained in the media until cells were harvested at 24 hours post-infection. ZIKV infection in DMSO treated cells elicited strong viral protein and RNA expression, high viral titers, and increased ATF4 and ATF3 levels - all consistent with ZIKV inducing the ISR pathway (Fig 3B-3G). However, in the presence of ISRIB, virus protein and RNA expression and virion production decreased (Fig 3B, 3F & 3G). The effects of ISRIB on ZIKV infection were not the result of inhibitor toxicity as a cell viability assay showed that treatment with 500 nM of ISRIB for 24 hours did not affect A549 cell growth (Fig 3H). These data show that the ISR pathway is an important modulator of ZIKV gene expression.

**FIG 3.**
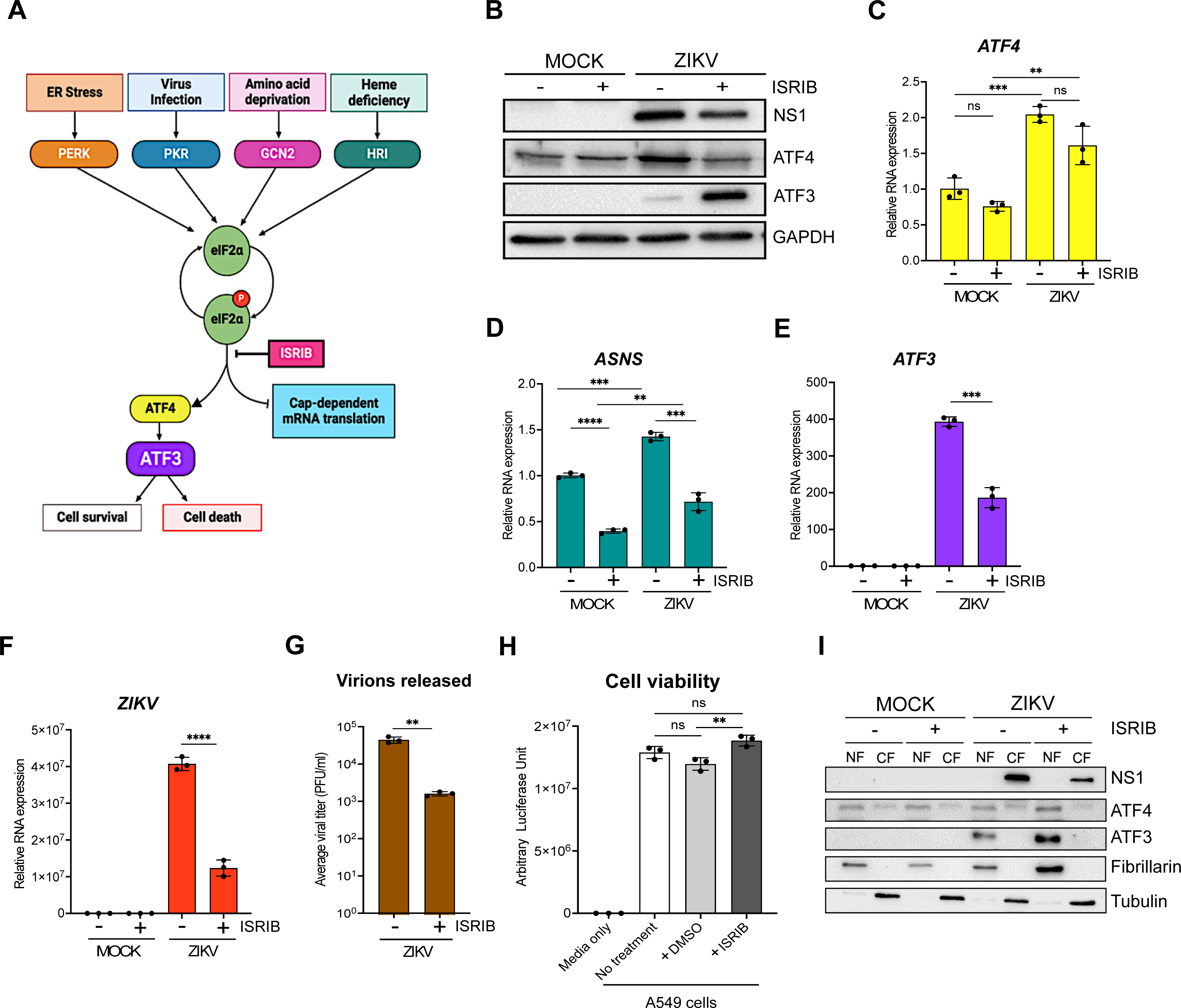
ZIKV activates ATF3 through the Integrated Stress Response (ISR) pathway. (**A**) Schematic of the ISR pathway. Stress conditions like virus infections, ER stress, amino acid deprivation and oxidative stress induce stalling of cap-dependent translation by phosphorylating eIF2⍺ and inducing the translation of ATF4. ATF4 in turn activates downstream targets including ATF3 to restore cellular homeostasis. A549 WT cells were mock-infected or infected with the ZIKV^PR^ (moi=10 PFU/cell) in the presence or absence of ISRIB, an ISR inhibitor. Cells were harvested 24-hours post-infection, and (**B**) cellular and viral proteins analyzed by western blot. The fold change (2^−ΔΔCt^) in (**C**) *ATF4*, (**D**) *ASNS*, (**E**) *ATF3* and (**F**) *ZIKV* mRNA levels relative to *ACTB* mRNA were determined by RT-qPCR from three technical replicates and three biological replicates. (**G**) Viral titers in cell culture media were measured by plaque assay. Average viral titers were calculated from three independent experiments with the paque assay being performed in duplicate. (**H**) A549 WT cells received no treatment or were incubated with DMSO (control) or ISRIB. Arbitrary luciferase unit were measured as a proxy for cell viability. (**I**) A549 cells were either mock-infected or infected with ZIKV^PR^ (moi=10 PFU/cell) in the presence or absence of ISRIB. Cellular and nuclear fractions were prepared from cells harvested 24-hours post-infection. The resultant subcellular fractions were analyzed by Western blotting and probed with anti-NS1, ATF4, ATF3, fibrillarin and α-tubulin antibodies. Fibrillarin and α-tubulin were used as nuclear and cytoplasmic markers respectively. The western bots shown are representative of three independent experiments. The quantitative data are shown as the means ± SD. The experiment was repeated three times. Statistical significance was determined by Student T-test. **p<0.05, ***p<0.0005, ****p<0.0001, ns-not significant. (18, 69)

We next examined the consequence of ISRIB on ATF4, the central integrator of the ISR pathway(20, 23). In mock-infected cells treated without or with ISRIB, ATF4 protein and RNA levels remained unchanged (Fig 3B & 3C). However, in ZIKV-infected ISRIB-treated cells ATF4 protein levels decreased and mirrored the levels in mock-infected cells in the absence or presence of ISRIB (Fig 3B). These data support the function of ISRIB as a pharmacological inhibitor of the ISR pathway. We also verified the inhibitor activity by measuring the mRNA levels of asparagine synthetase (*ASNS*), a well characterized downstream target that is transcriptionally controlled by ATF4 (57–59). Specifically in the presence of ISRIB, global cellular translation would progress in the absence or presence of a stressor. Consequently, ATF4 protein expression, and that of the downstream targets such as *ASNS*, would be suppressed (Fig 3B). Indeed, *ASNS* mRNA levels were reduced in both mock- and ZIKV-infected cells treated with ISRIB (Fig 3D). In contrast, ZIKV-infected cells treated with DMSO showed increased ATF4 protein and mRNA (Fig 3B & 3C), and increased *ASNS* mRNA abundance (Fig 3D).

Last, we examined ATF3 protein and mRNA expression (Fig 3B & 3E). ATF3 expression was not activated in mock-infected cells treated with DMSO or ISRIB. As expected, during ZIKV infection ATF3 mRNA and protein were expressed, while in the presence of ISRIB the levels of *ATF3* mRNA decreased (Fig 3E), consistent with the effect of ISRIB on ATF4 protein abundance (Fig 3B). Unexpectedly however, ATF3 protein levels notably increased with ISRIB treatment (Fig 3B). Since ATF3 is a transcription factor and functions in the nucleus, we next examined the subcellular localization of the increased protein levels. Here mock- and ZIKV-infected cells treated with DMSO or ISRIB were harvested, and the cytoplasmic and nuclear fractions isolated, and the protein distribution examined by western blot analysis (Fig 3I). We used fibrillarin and α-tubulin as cellular markers for the nuclear and cytoplasmic fractions, respectively. Subcellular fractionation showed that the increased levels of ATF3 protein in ZIKV-infected cells treated with ISRIB were present in the nuclear fraction (Fig 3I). These results show that following ZIKV infection and inhibition of the ISR pathway, the accumulated ATF3 predominantly localized to the nucleus.

Because *ATF3* mRNA levels decreased in ZIKV-infected cells treated with ISRIB but the protein significantly increased (Fig 3E & 3B), we examined whether this response was specific to the broad ISR inhibitor or if an ISR kinase-specific inhibitor would have the same response. We therefore treated mock- and ZIKV-infected cells without or with GSK2606414, an inhibitor that blocks autophosphorylation of PERK (60) and downstream activation of the ISR pathway induced by ER stress (Fig 3A). Like the effect of ISRIB, PERK inhibition decreased viral protein and RNA were expressed with ZIKV-infection (data not shown). ATF4 protein and mRNA levels on the other hand increased in ZIKV-infected cells treated with the PERK inhibitor (data not shown), which was likely the result of activation of the other ISR kinases (Fig 3A), such as PKR, in response to ZIKV-infection (52, 61). Notably in ZIKV-infected cells inhibition of PERK also decreased *ATF3* mRNA levels and increased ATF3 protein levels (data not shown). Overall, these results show that during ZIKV infection, ATF3 is activated through the ISR pathway, and is expected to modulate cellular stress by regulating transcription of specific genes. However, when the ISR pathway is inhibited, ATF3 protein expression may be upregulated, through either enhanced cap-dependent translation or mechanisms stabilizing the protein, which could control the cellular stress induced during viral infection. Future studies will examine the mechanism directing upregulation of ATF3 protein and downstream transcriptional control.

### ATF4 is the key activator of ATF3 during ZIKV infection

Our data show that the ISR pathway is an important regulator of ZIKV gene expression and contributor to ATF3 activation. Thus, we next investigated if ATF4, the master regulator of the ISR pathway, was the upstream activator of ATF3 during ZIKV infection. To this end, we depleted ATF4 with shRNAs stably transduced in A549 cells, and then either mock or ZIKV infected the A549 cells. As a control, we used A549 cells stably expressing a scramble non-targeting shRNA. Viral and cellular protein and RNA were analyzed 24 hours post-infection. To determine if depletion of ATF4 would affect ATF3 expression, we first treated cells with tunicamycin or DMSO (vehicle control) to induce ATF3 expression. In control non-targeting shRNA transduced cells treated with tunicamycin we observed an increase in ATF4 and ATF3 expression (Fig 4A & 4B). ZIKV infection upregulated ATF4 and ATF3 protein and RNA abundance (Fig 4A & 4B). Conversely, knock-down of ATF4 significantly reduced ATF3 levels in tunicamycin-treated and ZIKV-infected cells (Fig 4A & 4B). Interestingly, and in contrast to the deletion of ATF3 in A549 cells (Fig 2), we found that depletion of ATF4 decreased ZIKV protein and RNA levels (Fig 4A & 4C). These data suggest that in ZIKV-infected cells, ATF4 is the key activator of ATF3, and ATF4 expression acts to promote ZIKV gene expression.

**FIG 4.**
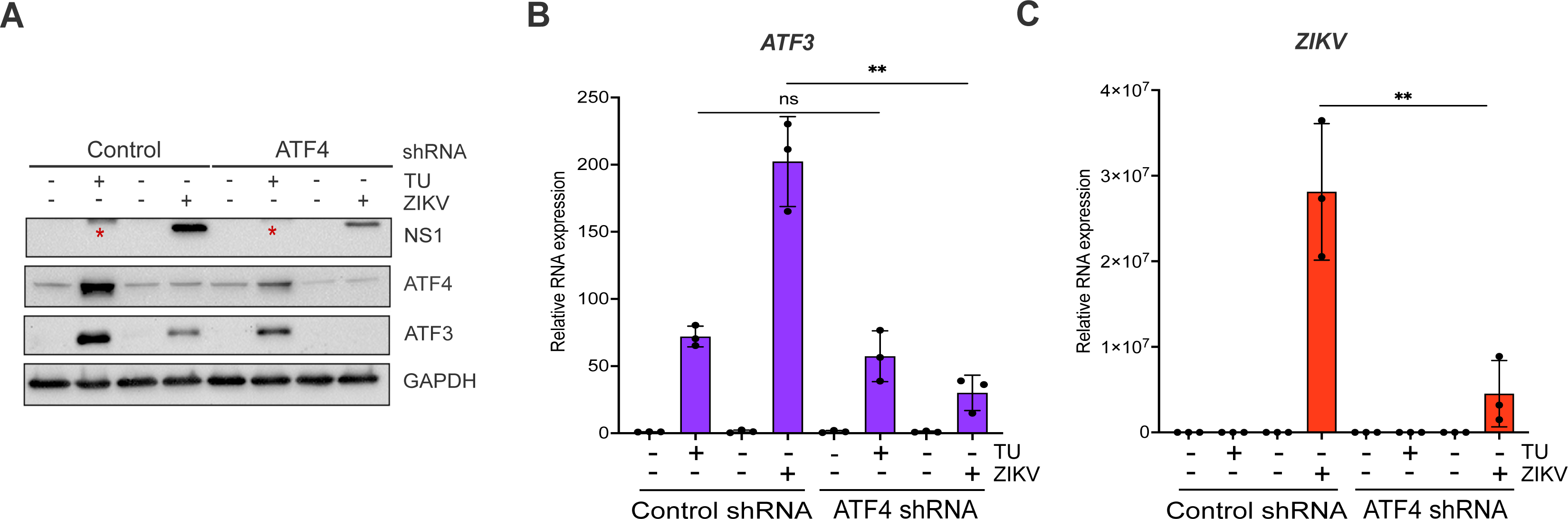
ATF4 induces ATF3 expression and promotes ZIKV protein and RNA expression. A549 WT cells stably expressing either control or ATF4 targeting shRNA were treated with tunicamycin (TU) or infected with ZIKV^PR^ (moi=10 PFU/cell). (**A**) ATF4, ATF3 and ZIKV NS1 proteins were assayed via western blot with GAPDH expression measured as the loading control. The blot shown is a representative of three separate experiments. (**B-C**) Fold change (2^−ΔΔCt^) in *ZIKV* and *ATF3* RNA expression relative to *ACTB* mRNA was determined by RT-qPCR. The RT-qPCR results presented are the mean ± SD of three technical replicates from three separate experiments. Statistical significance was determined by Student T-test. **p<0.05, ns-not significant, * non-specific band detected by the anti-NS1 antibody.

### ATF3 and ATF4 have opposing effects during ZIKV infection

ATF3 expression functions to restrict ZIKV gene expression, while the upstream effector protein ATF4 has a proviral role (Fig 2B, 2C, 2F, 2G, 4A & 4C). With these opposing functions, we hypothesized that if both ATF3 and ATF4 were depleted, viral expression would be restored to levels comparable with WT infected cells. To test this hypothesis, we transfected WT and ATF3 KO cell lines with either a control siRNA or siRNA targeting ATF4. These cells were then mock-infected or infected with ZIKV. By western blot and RT-qPCR we determined that ATF4 was successfully depleted in both WT and ATF3 KO cells (Fig 5A & 5B). Consistent with the data in Fig 4, depletion of ATF4 in WT cells decreased the abundance of ZIKV protein and RNA, and the expression of ATF3 (Fig 5A & 5C). In line with our prediction, we observed that ZIKV protein and RNA levels were rescued in cells lacking ATF3 and depleted of ATF4, albeit not to the same level as in WT A549 cells (Fig 5A & 5C). Therefore, ATF3 and ATF4 have opposing roles that together modulate the cellular response to ZIKV infection.

**FIG 5.**
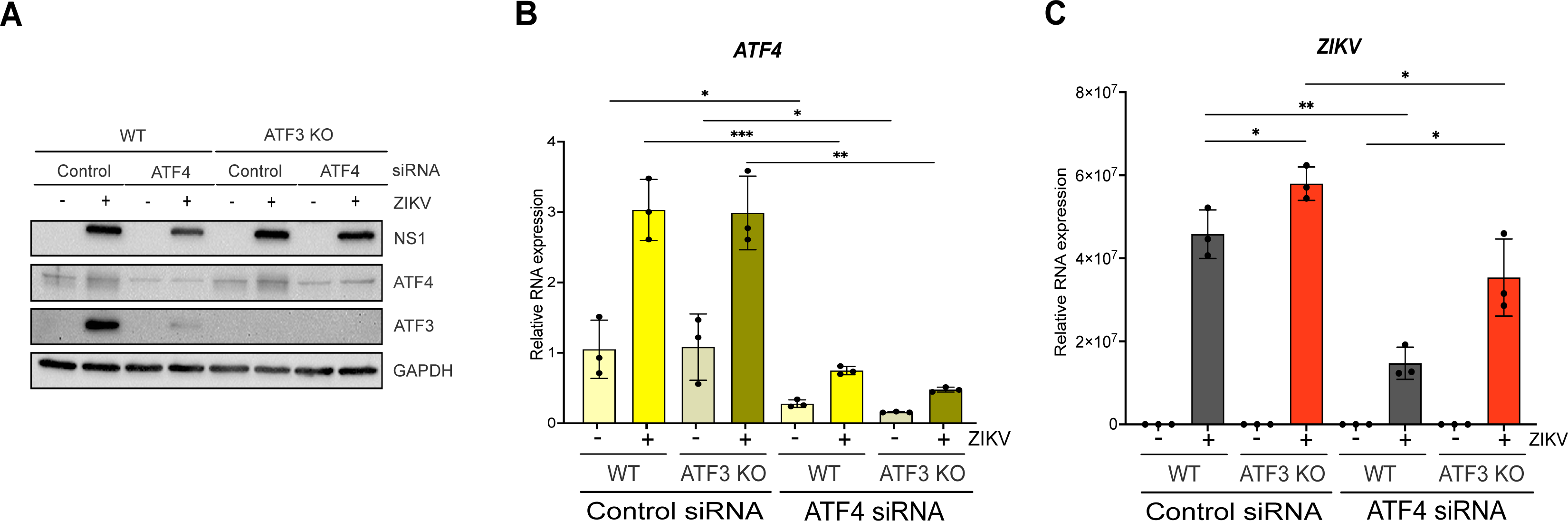
ATF3 suppresses while ATF4 promotes ZIKV RNA and protein expression. A549 WT and ATF3 KO cells expressing either control or ATF4 targeting siRNA were infected without or with ZIKV (moi=10 PFU/cell). (**A**) ZIKV NS1, ATF4 and ATF3 proteins were analyzed by western blot with GAPDH as the loading control. The western blot is a representative from three independent experiments. (**B-C**) Fold change (2^−ΔΔCt^) of *ATF4* and *ZIKV* RNA abundance relative to *ACTB* mRNA was determined by RT-qPCR. For each independent experiment, the RT-qPCR was performed in triplicate. N=3. The RT-qPCR data shown are the mean ± SD. Statistical significance was determined by Student t-test. *p < 0.05; **p < 0.01; ***p < 0.001.

### Global analysis of ATF3-dependent gene expression in response to ZIKV

In the absence of ATF3, ZIKV protein, RNA and titers increase (Fig 2). One mode by which ATF3 might restrict ZIKV gene expression is by regulating the transcription of distinct genes that antagonize ZIKV. To better understand the gene regulatory networks controlled by ATF3 that appear to restrict ZIKV infection, we compared changes in the polyA+ transcriptome of A549 WT and ATF3 KO cell lines after 24 hours of mock- or ZIKV-infection. Principal component analysis (PCA) revealed four clusters separating the samples by genotype (WT and ATF3 KO) and infection condition (mock and ZIKV) (Fig 6A). ZIKV infection induced substantial changes in the transcriptome in both, WT and ATF3 KO, genotypes. However, most transcripts had increased expression in both cell types after ZIKV infection with 1,769 transcripts being upregulated in WT and 2,184 transcripts being upregulated in ATF3 KO compared to the mock infection condition (Fig 6B & 6C). Upon closer investigation, more than half of upregulated transcripts in each genotype were shared (1,157), but a considerable number of transcripts were unique to each genotype (Fig 6D). These results suggest that ATF3 has a specific transcriptional role in the viral-induced stress response (Fig 6D).

**FIG 6.**
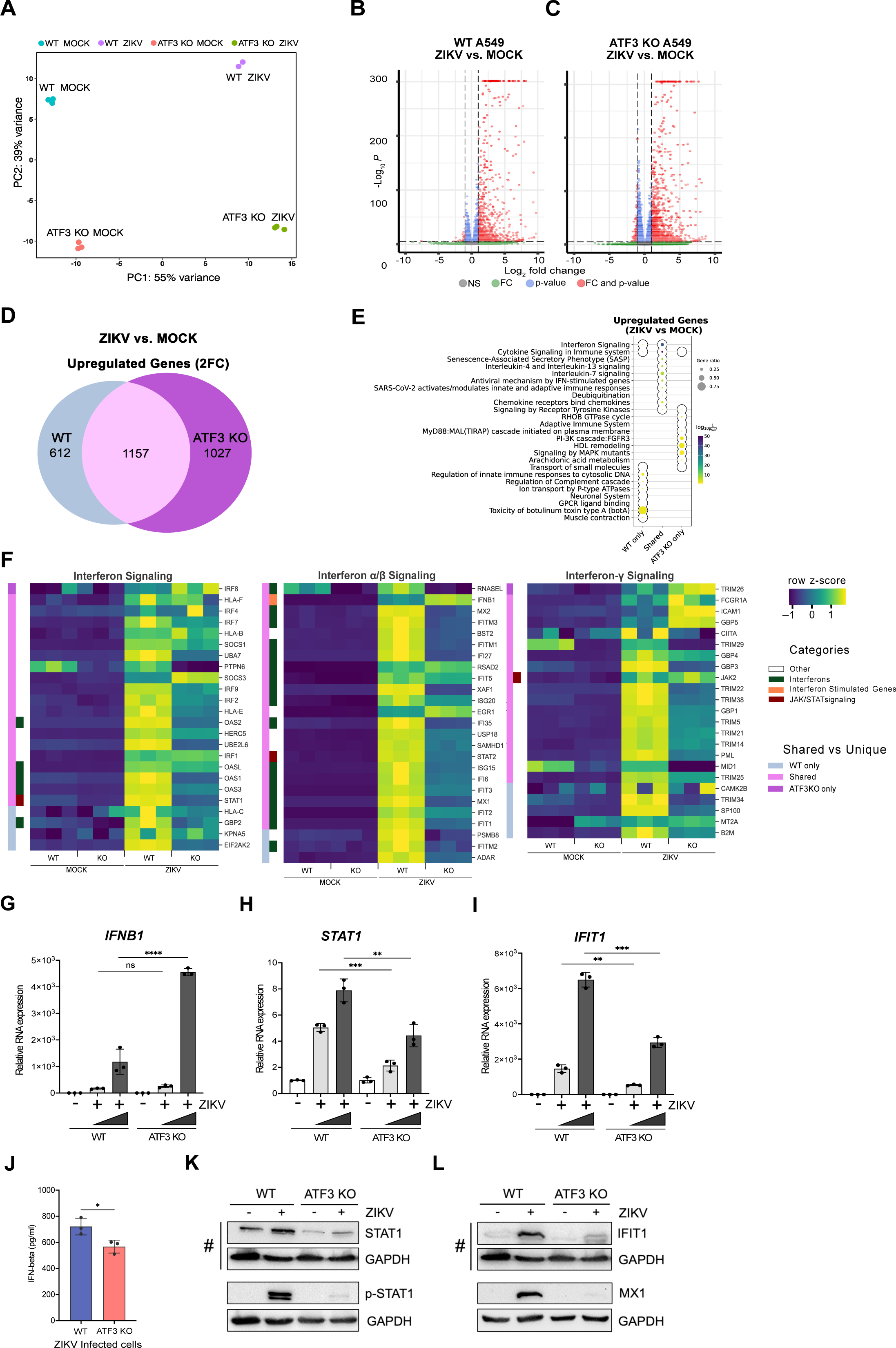
ATF3 regulates the antiviral immune response. A549 cells WT and ATF3 KO cells were mock-infected or infected with ZIKV^PR^ (moi=10 PFU/cell) and polyA-selected RNA expression was examined by RNA-seq analysis 24-hours post-infection. (**A**) PCA plot summarizing variance in gene expression in two A549 genotypes (WT and ATF3 KO) and infection conditions (mock and ZIKV). (**B**-**C**) Enhanced volcano plots showing differentially expressed genes in ZIKV-infected cells compared to (**B**) the mock-infected A549 WT or (**C**) ATF KO cells. The dotted lines represent the adjusted p-value threshold of 0.05 and fold-change (FC) threshold of 2. (**D**) The venn diagram shows the number of significantly (p-value < 0.05) upregulated (fold-change > 2; 2FC) genes in ZIKV-infected cells that are shared or unique between the two A549 cell genotypes (WT or ATF KO). (**E**) Metascape pathway enrichment analysis (Reactome) of shared and unique gene groups described in panel. The dotplot represents top ten enriched Reactome terms for each gene set. (**F**) Heatmaps showing relative expression values (row-normalized z-score) of genes associated with interferon signaling pathway that were significantly upregulated in ZIKV infection condition in at least one A549 genotype (WT or ATF3 KO). Genotype-specific shared or unique genes and select immune gene categories are highlighted in different colors as indicated in the legend. (**G-I**) mRNA expression of select innate immune response genes, (**G**) *IFNB1,* (**H**) *STAT1,* and (**I**) *IFIT1* were validated by RT-qPCR analyses following ZIKV infection (moi=1 and 10 PFU/cell). Target RNAs were normalized to *ACTB* mRNA, and the mRNA expression was determined by the 2^−ΔΔCt^ method. These RT-qPCR validation experiments were undertaken in three technical replicates from three separate experiments that were also independent from the RNA-seq samples. The data represent mean ± SD. Statistical significance was determined by Student T-test. *p<0.01, **p<0.001, ***p<0.0005, ****p<0.0001. (**J**) IFN-β protein secretion 24 hours post-infection was measured by ELISA in WT and ATF3 KO cells. The data show the mean ± SD. N=3. *p<0.05. (**K, L**) Protein expression of (**K**) STAT1 and phospho-STAT1 (p-STAT1), and (**L**) IFIT1 and MX1 were analyzed by western blotting. GAPDH levels were used as the loading control. # denotes the same western blot membrane that was probed for STAT1, IFIT1 and GAPDH. The images are separated into two panels, **K)** shows changes in STAT1 and phosphorylated STAT1, and **L)** shows the levels of IFIT1 and MX1 interferon induced proteins. Phosphorylated-STAT1 and GAPDH (and MX1 and GAPDH) proteins where blotted and probed on separate membranes. The blots shown are representatives from two separate experiments.

Next, we used gene set enrichment strategies to group differentially expressed genes into functional biological and phenotypic categories. We focused on the genes upregulated in response to ZIKV infection (Fig 6B & 6C). Pathway enrichment analysis suggested that most of the ZIKV-induced transcripts were immune response-associated genes (Fig 6E), in line with the expected cellular response to viral infection (62). More than half of the transcripts upregulated after ZIKV infection in WT cells were also significantly upregulated in ATF3 KO (Fig 6D), and these genes were primarily associated with interferon and cytokine signaling (Fig 6E). Despite significant upregulation in response to ZIKV relative to mock infection, many of the immune response-associated genes displayed dampened induction and lower overall transcript abundance in ATF3 KO relative to WT (Fig 6F). Select genes involved in interferon signaling and innate immune responses were induced by ZIKV only in WT cells (Fig 6F). ZIKV-induced genes specific to ATF3 KO were associated with cellular metabolism and cell structural components like membrane lipids and cytoskeletal components (Fig 6E). The impact of these ATF3 KO-specific ZIKV targets could, for example, affect the formation of ZIKV replication complexes, regulation of autophagy, and ZIKV pathogenesis (19, 63–65). Overall, these global gene expression data are consistent with the hypothesis that ATF3 positively regulates the transcription of genes involved in the innate immune response as one mechanism to restrict ZIKV infection.

To validate the RNA-seq data, key genes involved in the IFN-induced antiviral pathway including *IFNB1*, *STAT1*, *IFIT1, MX1, ISG15, IRF9, OASL,* and *DDX58/RIG-I* were chosen for RT-qPCR (Fig 6G-I, and data not shown) and ELISA or immunoblot analyses (Fig 6J-M). We analyzed mRNA expression in WT and ATF3 KO cells that were mock- or ZIKV infected. From our RT-qPCR results, the expression pattern of all genes tested reflected the expression profiles from our RNA-seq analysis (Fig 6F-I). At the protein level, the secreted IFN-β protein, as measured by ELISA was significantly lower in ATF3 KO ZIKV-infected cells despite the RNA levels being higher (Fig 6G & 6J). This decrease in the amount of secreted IFN-β might be a consequence of translational regulation, ER stress, and effects on vesicular trafficking. By immunoblot, the protein levels of STAT1 in ATF3 KO cells, without and with ZIKV infection, were notably decreased compared to WT cells (Fig 6K). The decrease in STAT1 protein levels affected the levels of STAT1 phosphorylation (Fig 6K) and in turn the abundance the interferon stimulated IFIT1 and MX1 mRNAs and proteins (Fig 6I & 6L, and data not shown). Alternatively, the absence of ATF3 might also transcriptionally affect IFIT1 and MX1 mRNA, and protein levels. Altogether, these data indicate that ATF3, either directly or indirectly, enhances the expression of antiviral immune response genes during ZIKV infection.

### ATF3-mediated antiviral immune enhancement is specific to ZIKV-infection

Poly I:C, a synthetic double-stranded RNA mimic, can activate double-stranded (ds) RNA sensors such TLR3 in the endosome and RIG-1 and MDA-5 in the cytoplasm (66–68). Induction of these sensors converge on IRF3 resulting in IFN-α/β expression (62, 69). To determine if the role of ATF3 in enhancing the antiviral response was specific to ZIKV, we examined the levels of select IFN-stimulated antiviral genes post poly I:C transfection in WT and ATF3 KO A549 cell lines (Fig 7A-D). Poly I:C induced expression of *STAT1*, *IFIT1*, and *MX1* in WT cells (Fig 7A-C). Notably, following poly I:C transfection the transcript levels of *STAT1*, *IFIT1*, and *MX1* further increased in ATF3 KO cells compared with WT cells, (Figures 7A-C). Protein analysis showed a modest increase in STAT1 but not STAT2 proteins, and the presence of phosphorylated STAT1 and STAT2 and induced IFIT1, MX1, and ATF3 proteins in WT in transfected with poly I:C (Fig 7D). In ATF3 KO cells, the levels of STAT1 but not STAT2, were reduced compared to WT cells, and poly I:C had no effect on these proteins (Fig 7D). Poly I:C treatment in ATF3 KO cells, like WT cells, induced phosphorylation of STAT1 and STAT2 (Fig 7D). Notably, the protein levels of IFIT1 and MX1, consistent with the mRNA levels, were higher in ATF3 KO cells than in WT cells (Fig 7B-D). Unlike our observation after ZIKV infection, ATF3 in WT cells in response to dsRNA mimic poly I:C negatively affects the expression of IFN response genes.

**FIG 7.**
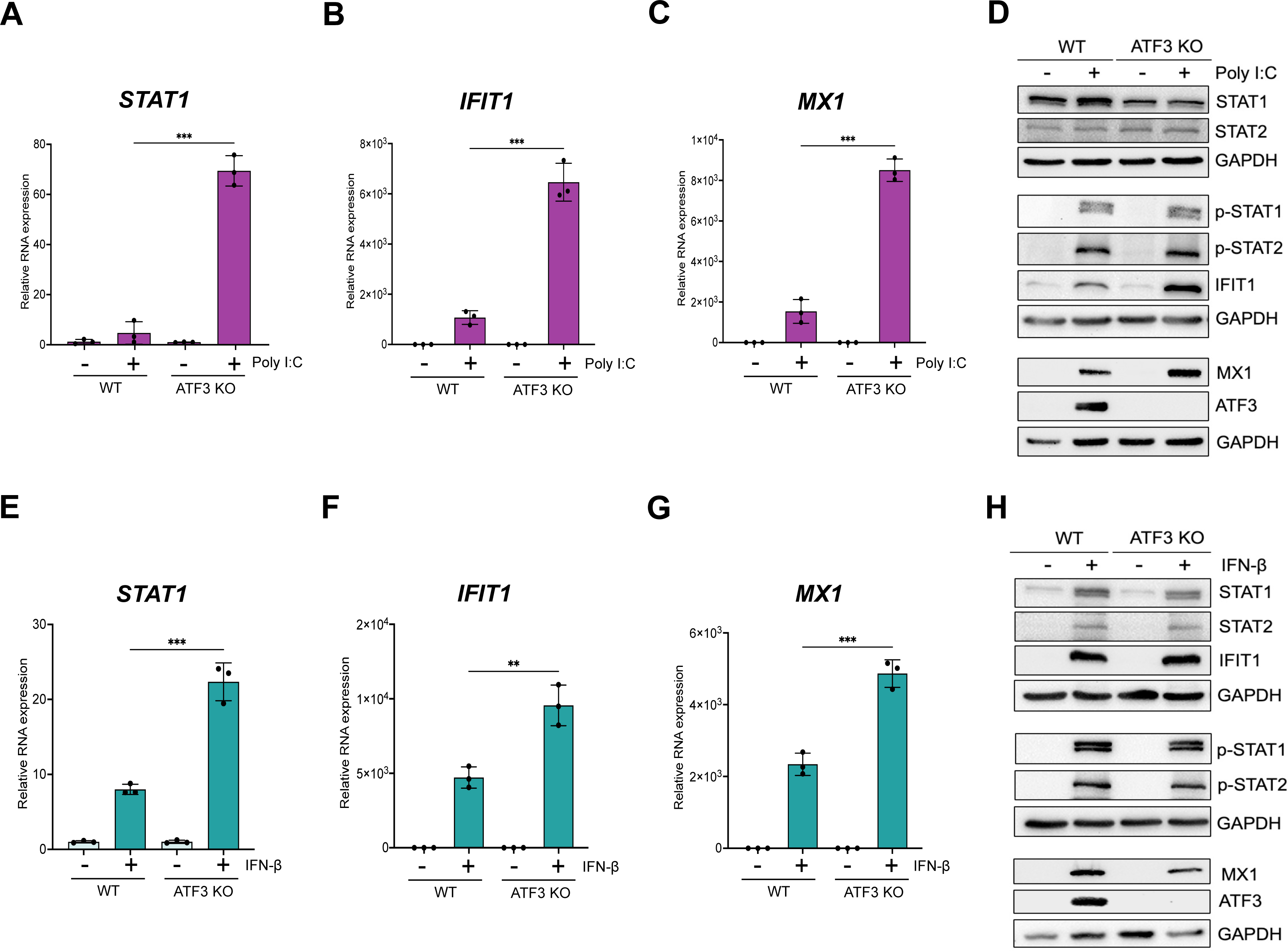
ATF3 selectively represses the expression of factors involved in IFN signaling upon poly I:C or IFN-β stimulation. A549 WT and ATF3 KO cells were transfected with 1 μg/ml poly I:C for 6 hours at 37°C. Total RNA was analyzed by RT-qPCR and relative mRNA expression was measured for (**A**) *STAT1*, (**B**) *IFIT1*, and (**C**) *MX1*. Target mRNAs were normalized to *ACTB* mRNA and relative transcript expression was calculated using the fold change (2^−ΔΔCt^) method. The results presented are means ± SD of three technical replicates from three independent experiments. (**D**) Western blot analysis shows the abundance of proteins associated with the antiviral immune response following poly I:C transfection. The experiment was repeated three times, and a representative blot is shown. A549 WT and ATF3 KO cell lines were treated with 10 ng/ml IFN-β for 24 hours at 37°C. Total RNA was isolated for RT-qPCR analysis using primers specific for (**E**) *STAT1*, (**F**) *IFIT1* and (**G**) *MX1*. The relative expression of each transcript was determined as described above. The data shown are the mean ± SD of three technical replicates of three independent experiments. (**H**) Cell lysates from IFN-β treated WT and ATF3 KO cells were used to analyze by antiviral (STAT1, STAT2, p-STAT1, p-STAT2, IFIT1, MX1) and ATF3 protein expression by western blotting. GAPDH was used as the loading control. A representative blot from three independent experiments is shown. Statistical significance of the RT-qPCR data was determined by Student T-test. **p<0.05, ***p<0.001.

To further validate the specific regulation of ATF3 observed with ZIKV, we exposed both cell lines to IFN-β, which activates the JAK/STAT signaling cascade to initiate the type-1 IFN antiviral pathway and production of interferon stimulated genes. In response to IFN-β treatment, RT-qPCR analysis showed an increased expression of *STAT1*, *IFIT1,* and *MX1* in WT cells (Fig 7F-G). Additionally, the expression of these genes was significantly higher in the ATF3 KO cells (Fig 7F-G). Following incubation with IFN-β of WT cells, we show by immunoblot that STAT1 and STAT2 were phosphorylated, and downstream IFN-stimulated IFIT1 and MX1 were expressed, indicating that the JAK/STAT signaling pathway was activated (Fig 7H). IFN-β treatment of WT cells also induced ATF3 expression (Fig 7H). Notably, in cells lacking ATF3 the abundance of STAT1 and MX1 were decreased, (Fig 7H) even though the mRNA transcripts were elevated (Fig 7E & 7G) and STAT1 was robustly phosphorylated (Fig 7H). Despite the increased of *IFIT1* mRNA levels in ATF3 KO cells following incubation with IFN-β (Fig 7F), IFIT1 protein levels were only modestly increased (Fig 7H). Overall, these data show that the innate immune response pathway when activated by either a synthetic double-stranded RNA mimic or following IFN-β treatment is not hindered by the absence of ATF3. Moreover, ATF3 restricts the expression of select transcripts within the type-1 IFN pathway under these conditions.

### ATF3 acts on genes within the JAK/STAT pathway to limit ZIKV infection

In response to viral infection, the innate immune pathway is activated to restrict virus infection (62, 69). In particular, the primary response is initiated by pattern recognition receptors which recognize different viral components and lead to expression of type 1 interferons (e.g., IFN-β) (62, 69). The release of interferon initiates the secondary innate immune response and expression of interferon stimulated genes (ISGs) which establish an antiviral state within the cell (62, 69). Our data indicate that ATF3 promotes the expression of components within the innate immune response pathway to restrict ZIKV infection (Fig 6). To investigate whether ATF3 affects ZIKV gene expression when the innate immune response is blocked, we selectively inhibited JAK1 and JAK2, key tyrosine kinases in the JAK/STAT signaling pathway using Ruxolitinib (70, 71) and infected WT and ATF3 KO cells for 24 hours. In WT cells, Ruxolitinib treatment inhibited the phosphorylation of STAT1 in response to ZIKV infection (Fig 8D), which blocked the expression of downstream ISGs such as IFIT1, MX1, OASL, and ISG15 (Fig 8B-D, and data not shown) and increased the abundance of ZIKV RNA to levels similar to ZIKV infection in ATF3 KO cells (Fig 8A). In ZIKV-infected ATF3 KO cells, Ruxolitinib similarly inhibited the JAK-STAT signaling pathway to restrict downstream IFN stimulated responses (Fig 8B-D). Moreover, following Ruxolitinib treatment the abundance of *MX1* mRNA was not significantly different between WT and ATF3 KO ZIKV-infected cells (Fig 8C). In contrast, *IFIT1* mRNA levels were elevated in ZIKV-infected ATF3 KO cells treated with Ruxolitinib compared to WT cells (Fig 8B) but this modest increase did not result in detectable IFIT1 protein (Fig 8D). Notably, ZIKV RNA levels were similar in ATF3 KO cells in the absence or presence of Ruxolitinib (Fig 8A). These data show that ATF3 expression affects components within the JAK/STAT signaling cascade to suppress ZIKV gene expression and virion production. In particular, the decreased expression of STAT1 in ATF3 KO cells (Fig 6 and Fig 8D), could be the central component which attenuates the downstream IFN stimulated response. ATF3 was previously shown to bind the STAT1 promoter region in murine cells (37), which presents the possibility that in A549 cells STAT1 is similarly transcriptionally controlled by ATF3, although such interactions remain to be determined.

**FIG 8.**
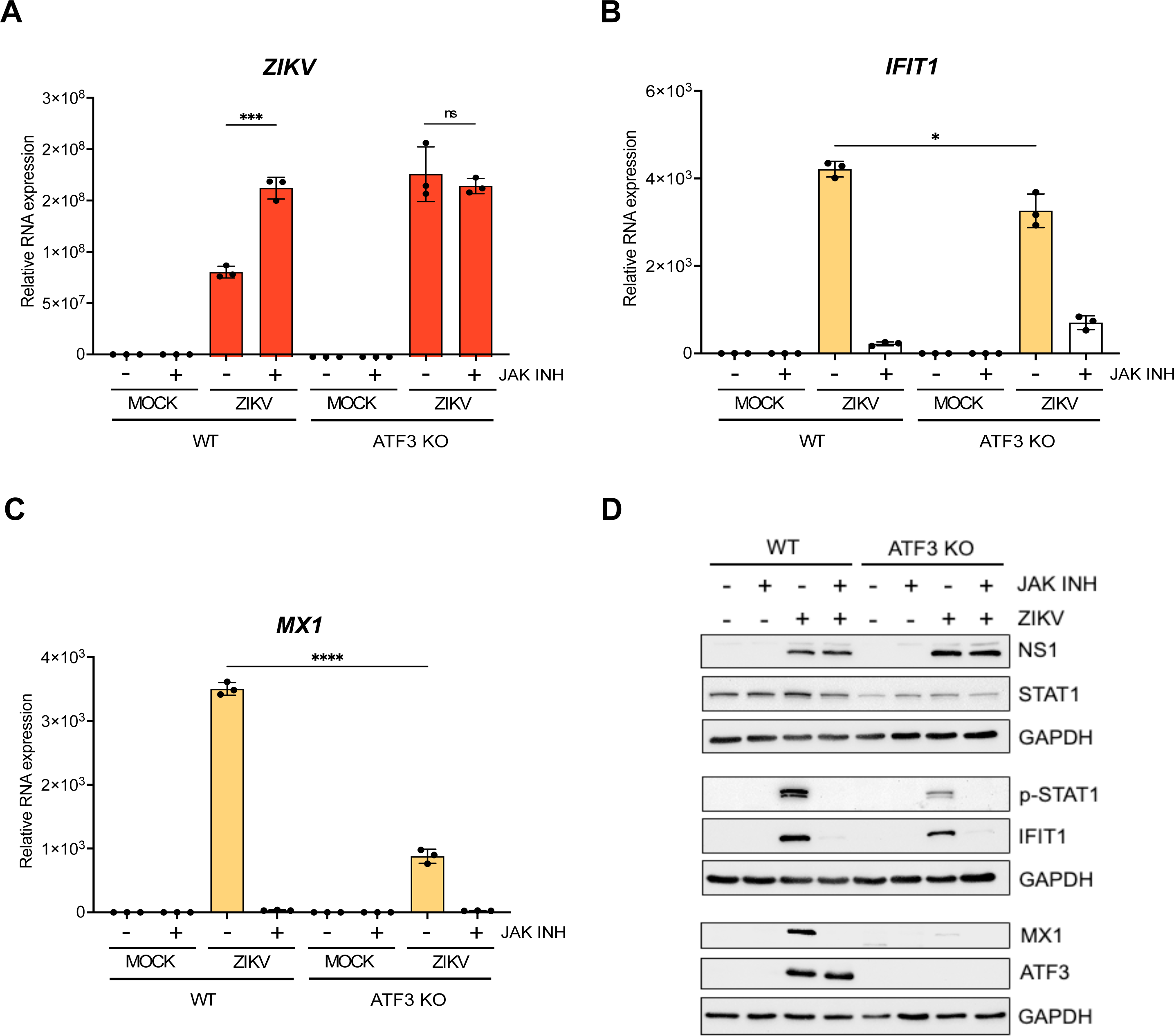
ATF3 restricts ZIKV infection through regulation of components within the JAK/STAT antiviral response pathway. A549 WT and ATF3 KO cells were mock- or ZIKV^PR^-infected (moi=10 PFU/cell) and co-treated with 30 nM of the JAK1/2 inhibitor Ruxolitinib for 24 hours at 37°C. Total RNA isolated from cells was used as a template for RT-qPCR analysis. (**A**) *ZIKV*, (**B**) *IFIT1* and (**C**) *MX1* RNA expression relative to *ACTB* were determined by 2^−ΔΔCt^ RT-qPCR method. The experiments were repeated three times and the data shown are the mean ± SD of triplicate measurements. Statistical significance was determined by Student T-test. *p<0.05, ***p<0.001, ****p<0.0001, ns-not significant. (**D**) A representative Western bot showing the expression of ZIKV NS1, antiviral (STAT1, p-STAT1, IFIT1 and MX1) and ATF3 proteins. N=3.

## Discussion

ATF3 mediates adaptive responses via the positive or negative modulation of cellular processes including immune response, autophagy, and apoptosis (31–33). For virus infections, ATF3 expression can produce anti-viral outcomes by regulating the transcription of host antiviral genes (38, 72, 73). Conversely this stress-induced transcription factor may benefit the virus by dampening the expression of genes necessary for virus restriction and/or resolution of virus-induced stress (36–38, 74). We previously showed that ATF3 was upregulated during ZIKV infection of SH-SY5Y cells and PMBC isolated from early acute ZIKV-infected patients (39, 40), however the upstream effector proteins inducing ATF3 expression and the impact of ATF3 activation on ZIKV gene expression was unknown.

In this study we determined that peak ATF3 expression coincides with robust ZIKV protein and RNA expression at 24 hours after infection in A549 cells (Fig 1). We identified the ISR pathway as the upstream signaling cascade leading to ATF3 activation during ZIKV infection (Fig 3) with ATF4 as the direct effector of ATF3 in this pathway (Fig 4). This observation is consistent with ZIKV activating the ISR through the ER sensor PERK and double-strand RNA sensor PKR (41, 51, 53). Upon stress induction, these kinases phosphorylate eIF2⍺ leading to the attenuation of global protein synthesis. This event initiates ATF4 translation and subsequently, ATF4 induces ATF3 expression (20, 75). Finally, we show that ATF3 enhances the expression of innate immune response (Fig 6) to suppress ZIKV gene expression (Fig 2). Overall, these data reveal important crosstalk between the integrated stress response pathway, ATF3 and antiviral responses during ZIKV infection.

Virus activation of the ISR either protects the against viral infections or is subverted by the virus to promote viral replication. Evidence of these roles has been demonstrated in several studies involving viruses within the *Flavivirus genus* (76–81). For example, in a JEV infection model, the JEV NS2A protein counteracted the antiviral effects of the ISR by specifically blocking PKR activation and eIF2⍺ phosphorylation, thereby ensuring effective viral replication (79). Similarly, during DENV infections in Huh7 and A549 cells, stimulation of PERK and IRE-1⍺ signaling led to increased viral replication (80). However, in the case of West Nile virus (WNV), previous reports indicated that infection induced PERK and PKR kinases lead to apoptosis and repressed viral replication (76, 81). Like other flaviviruses, ZIKV infection activates the PERK arm of the ISR pathway in human neural stem cells, and in embryonic mouse cortices after intra-cerebroventricular injection with the virus (51). This activation of the ISR pathway increased *ATF4*, *ATF3* and *CHOP* mRNA levels and caused a disruption in the proper formation and survival of neurons during cortical development. Interestingly, co-treatment with the PERK inhibitor GSK2656157 attenuated this outcome (51). Consistent with these data, in A549 ZIKV-infected cells, we observed that GSK2656157 inhibited the phosphorylation and activation of PERK and ATF4 translation, which reduced *ATF3* and *PERK* mRNA accumulation and decreased ZIKV protein and RNA levels (data not shown).

When we inhibited the ISR pathway during ZIKV infection using ISRIB, a broad ISR inhibitor (Fig 3) or GSK2606414, a PERK inhibitor (data not shown), ATF4 protein expression was reduced and *ATF3* mRNA levels were negligible (Fig 3, and data not shown). These results align with ATF4 being the upstream effector protein of ATF3 in the ISR pathway (Fig 4). Unexpectedly however, ATF3 protein, but not the mRNA, levels dramatically increased (Fig 3) following inhibition of the ISR and ZIKV infection. Of note, we did not see this same response following tunicamycin treatment and inhibition of the PERK pathway (data not shown). Consistent with the transcriptional role of ATF3, we observed that the protein was predominantly in the nucleus (Fig 3I). At present, the mechanism leading to increased ATF3 protein levels is unknown. One possibility might be that following ISRIB treatment and ZIKV infection, the low levels of *ATF3* mRNA are more efficiently translated via the cap-dependent mechanism. We also considered that, like ATF4, ATF3 might be translationally regulated via an upstream open reading frame (28). However, inspection of the 5’ UTR revealed a short UTR length and the absence of an upstream (or downstream) AUG codon that could direct this stress-induced translational control mechanism. Alternatively, under the appropriate stress conditions, ATF3 protein levels could be regulated by either an alternate translational control mechanism such as via an internal ribosomal entry site and/or protein stability/turnover pathway (82, 83). Indeed, ATF3 protein stability has been shown to be regulated by UBR1/UBR2 and MDM2 ubiquitinases, and the ubiquitin-specific peptidase 33 (USP33) protein (84, 85). It is therefore possible that differential expression of ubiquitinases and/or deubiquitinases during ZIKV infection and inhibition of the ISR pathway changed ATF3 protein levels. Additional experiments are however needed to investigate such regulation.

ATF4 is a master regulator of the ISR pathway (20, 23, 86). During ZIKV infection we observed increased levels of ATF4 RNA and protein (Fig 1A-B), and this increase in ATF4 expression led to the activation of ATF3 (Fig 4). This result aligns with a previous study showing that ATF3 is regulated redundantly by two different stress-dependent pathways: the ATF4-dependent ISR pathway and the p53 gene regulatory network (87). Specifically, ATF4 regulates ATF3 directly at the transcript level via promoter binding and regulation but if ATF4 is inhibited or depleted, ATF3 can still be turned on by other pathways (87). In contrast to the antiviral effects of ATF3, we determined that depleting ATF4, led to a decrease in ZIKV protein and RNA expression (Fig 4). Proviral functions of ATF4, such as directly controlling cellular transcription to promote human immunodeficiency virus 1 (HIV-1), human herpes virus 8 (HHV-8), and murine cytomegalovirus (MCMV) infections, have been described (88–92). While the mode by which ATF4 positively regulates ZIKV remains to be determined, one possibility could be the activation of ATF4-dependent genes like GADD34 (growth arrest and DNA damage-inducible protein 34) which downregulate the ISR by recruiting protein phosphatase 1 (PP1) to dephosphorylate eIF2⍺, promote ZIKV translation and downstream steps in the infectious cycle (41, 93). ATF4 was also found to positively affect porcine reproductive and respiratory syndrome virus (PRRSV), a single-stranded positive-sense RNA virus that replicates in cytoplasm (41). Thus, like PRRSV, ATF4 could be affecting a specific step(s) in the ZIKV infectious cycle. Regardless, future studies are needed to uncover the mode by which ATF4 positively regulates ZIKV.

ATF3 affects a host of cellular systems, including cell cycle (94), apoptosis (95), neuron regeneration (96, 97), serine and nucleotide biosynthesis (98, 99) and the immune response (31). For the latter, ATF3 functions have been described as a rheostat that regulates the immune response (31). For instance, in ATF3-deficient bone marrow-derived macrophages (BMDM), the expression of IFN-β and other downstream components were upregulated compared to WT cells, and this attenuated LMCV and VSV*DG(Luc) replicon infections (74). In NK cells, ATF3 negatively regulated IFN-γ expression however, the reverse was observed in MCMV infected ATF3 knockout mice compared to WT mice (38). Similarly, ISGs were upregulated in JEV infected Neuro2A and MEF cells depleted of ATF3, and chromatin immunoprecipitation studies showed that ATF3 bound to select promoter regions in STAT1, IRF9, and ISG15 (37). Given these prior studies showing ATF3 regulating the immune response, we reasoned that ATF3 transcriptionally controlled genes involved in the innate immune response, to promote ISG expression and restrict ZIKV infection. From our RNA-seq data, the absence of ATF3 specifically led to a decrease in the transcription of genes involved in IFN pathways (Fig 6) which supports the role of ATF3 as a positive transcriptional regulator of these genes during ZIKV infection. Notably, depletion of ATF3 did not suppress all innate immune effectors as *IFNB1* (IFN-β) was upregulated in both WT and ATF3 KO cells (Fig 6F & 6G), and *IFNB1* mRNA levels were significantly higher in the ATF3 KO cells compared to WT cells (Fig 6G). In BMDM, two ATF3 binding sites were identified in the promoter and upstream region of *IFNB1,* where the second binding site functioned to negatively regulate *IFNB1* levels (74). It is possible that in our A549 KO system this second binding site is nonfunctional and thus *IFNB1* expression is not subjected to feedback regulation. Alternatively, other studies predict that ATF3 potentially suppresses interferon expression by remodeling nucleosomes and keeping chromatin in a transcriptionally inactive state through interacting with histone deacetylase 1 (74, 100). Future transcriptomic studies defining ATF3 genomic occupancy during ZIKV infection will elucidate how this stress induced transcription factor differentially directs the expression of *IFNB1* and other ISGs. Last, the higher abundance of *IFNB1* in ATF3 KO cells did not result in increased IFN-β. Instead, the amount of IFN-β secreted from the ATF3 KO cells was less than in WT cells (Fig 6J). One possibility for this difference could be that with increased levels of ZIKV infection (Fig 2), ER stress may persist which would affect the overall trafficking of secreted proteins such as IFN-β.

Sood and colleagues first showed that ATF3 was upregulated during JEV infection and that RNAi depletion of ATF3 decreased JEV protein and RNA abundances as well as viral titers (101). Moreover, during JEV infection, ATF3 was reported to negatively regulate antiviral response and autophagy, likely by controlling transcription (37). In contrast, our findings indicate that ATF3 functions as a positive effector of the antiviral response (Fig 6), specifically targeting genes within the type-1 IFN pathway to suppress ZIKV gene expression and virion production (Fig 8). These differences might be explained by differences in the cell types used in these experiments and/or impact of dimerization on ATF3 function. ATF3 can have both activator and repressor functions (34, 102), depending on whether this stress inducible transcription factor homodimerizes or forms a heterodimer with other transcription factors. The previous JEV studies were conducted using mouse Neuro2A and mouse embryonic fibroblast cells (37), while we used human A549 lung adenocarcinoma and HCT-116 colorectal carcinoma cells (Fig 2). Differences in the abundance of interacting partners between mouse and human cell lines may influence ATF3 dimerization and thus the transcriptional responses. Alternatively, as JEV and ZIKV belong to different flavivirus clades, the difference in ATF3 function may be related to a virus specific response. Future studies are needed to elucidate the virus genetic determinants that modulate ATF3 function.

Finally, we investigated how ATF3 might enhance the interferon response during ZIKV infection. To this end we treated WT and ATF3 KO cells with either poly I:C or IFN-β or inhibited the JAK/STAT signaling pathway with Ruxolitinib. In contrast to ZIKV-infection in ATF3 KO cells, poly I:C and IFN-β treatment of ATF3 KO cells activated JAK/STAT signaling, increased the transcript levels of *STAT1*, *IFIT1,* and *MX1* (Fig 7A-C & Fig 7E-G), and led to the expression of downstream IFN-induced proteins (Fig 7D & 7H). These data showed that the ATF3 effect on the innate immune response is ZIKV-specific. These data also highlight the ability of ATF3 to discern various stressors, enabling context-specific regulation consistent with the role as a transcriptional regulator (34, 49). The difference in ATF3 function under poly I:C or IFN-β treatment conditions may stem from differences in upstream pathways activating ATF3 or the selection of binding partners that regulate the transcription of downstream targets (31, 33). Future studies addressing these questions will provide mechanistic insights into the impact of diverse stimuli on ATF3 activation and downstream regulatory effects particularly on the interferon response.

When we investigated the effect of ATF3 expression on the JAK/STAT pathway by treating ZIKV-infected cells with Ruxolitinib, the JAK1 and JAK2 inhibitor, viral RNA expression was predictably increased compared to control-treated WT cells (Fig 8A). Interestingly, ATF3 depletion alone, or in combination with JAK inhibition led to an increase in viral RNA levels similar to ZIKV-infected WT cells treated with Ruxolitinib (Fig 8A). These data suggest that ATF3 targets the JAK/STAT pathway to enhance antiviral response to ZIKV. With ATF3 binding sites previously identified in the promoter regions of STAT1 in mouse neuronal cells (37) and ATF3 recently shown to promote STAT1 expression in a diabetic injury model (103), we posit that, ATF3 directly regulates STAT1 transcription within the JAK/STAT pathway to enhance the antiviral response against ZIKV infection (Fig 2 & Fig 6). By regulating STAT1 abundance in response to ZIKV-infection, downstream effects following IFN-β (and/or IFN-γ) induction of the pathway (Fig 6F) would impact ISG expression and functions. Future studies that establish the direct targets of ATF3 particularly within the IFN pathway and the type of regulation will provide valuable mechanistic insights on the role of ATF3 during ZIKV infection.

In summary, our study demonstrates that during ZIKV infection, the stress-induced transcription factor ATF3, activated through the ISR pathway and ATF4, enhances antiviral response by directly influencing the expression of genes involved in the JAK/STAT signaling pathway and regulation of the antiviral state. Our findings reveal important crosstalk between the ISR and antiviral response pathway through ATF3. Overall, our work contributes to a deeper understanding of the complex interplay between ZIKV infection, cellular stress pathways, and transcriptional control and the impact on infection outcomes.

## Materials and Methods

### Cell Lines and ZIKV

A549 (Human lung epithelial adenocarcinoma, ATCC CCL-185) wild type (WT) and ATF3 knock-out (KO) cell lines were maintained in Dulbecco’s minimal essential medium (DMEM; Gibco, #11995-065) supplemented with 10% fetal bovine serum (FBS; Seradigm, #97068-085), 10 mM nonessential amino acids (NEAA; Gibco, #11140076), 2 mM L-glutamine (Gibco, #25030081) and 1mM sodium pyruvate (Gibco, #11360070). The HCT-116 wild-type and ATF3 knockout cell lines were generously provided by Dr. Chunhong Yan, Augusta University (49). These cells were grown in McCoy’s 5A media (Corning, #10-050-CV) supplemented with 10% FBS (Seradigm, #97068-085) and 1% penicillin and streptomycin (Gibco, #15140163). Vero cells (ATCC CRL-81) were cultured in DMEM (Gibco, #11995-065) supplemented with 10% FBS (Seradigm, #97068-085), 1% penicillin and streptomycin (Gibco, #15140163) and 10 mM HEPES (Gibco, #15630080). HEK 293FT cells (Invitrogen, #R70007) were grown in DMEM (Gibco, #11995-065) with 10% FBS (Seradigm, #97068-085), 10 mM NEAA (Gibco, #11140076) and 2 mM L-glutamine (Gibco, #25030081). All cell lines were cultured at 37°C with 5% CO_2_ in a water-jacketed incubator. ZIKV^PR^ (Puerto Rico PRVABC59) strain was a gift from Dr. Laura Kramer (Wadsworth Center NYDOH) with permission from the CDC. Viral stocks were prepared in C6/36 cells (ATCC CRL-1660) by infecting near confluent cells at a multiplicity of infection (moi) of 0.1 and incubating at 28°C. At 7 days post-infection, media from infected cells were collected and aliquots supplemented with 20% FBS were stored at -80°C. Viral RNA was extracted and examined by RT-qPCR and viral titers were measured by plaque assay to validate infection.

### Creating the ATF3 Knock-out (KO) A549 Cell Line

We generated A549 ATF3 KO cells in our laboratory using the CRISPR/Cas9 system. The following gRNA sequence targeting ATF3 was cloned into pLentiCRISPRv2 plasmid: 5’-CCACCGGATGTCCTCTGCGC-3’ (Genscript, Clone ID C88007). HEK 293FT cells were co-transfected with pLentiCRISPRv2-ATF3 CRISPR gRNA, and pMD2.G (Addgene, #12259) and psPAX2 (Addgene, #12260) packaging plasmids using JetOptimus DNA transfection reagent (Polyplus, #101000025) according to the manufacturer’s protocol. Media containing lentivirus was collected 24- and 48-hours post-transfection and pooled together. The pooled lentivirus media was filtered through a 0.45 mm pore filter and used to transduce A549 cells in the presence of 6 μg/ml polybrene (Sigma-Aldrich, TR1003). Twenty-four hours later, the lentivirus-containing media was removed, replaced with fresh media and cells were incubated at 37°C. After 24 hours of incubation, the transduced cells were transferred into new tissue culture dishes and puromycin (1 μg/ml) (InvivoGen, #ant-pr-1) selection was carried out for 4 days by which time all A549 WT control cells were killed by the antibiotic. Individual clones were isolated by diluting, seeding in a 96-well plate, and incubating at 37°C. Following expansion, clones were screened in the absence and presence of tunicamycin and ATF3 expression determined by western blotting and RT-qPCR. DNA was also isolated from successful KO clones using DNAzol (Invitrogen, #10503027) reagent. PCR was subsequently carried out with forward and reverse primers (5’-CTGCCTCGGAAGTGAGTGCT-3’ and 5’-AACAGCCCCCTGCCTAGAAC-3’) that spanned part of the *ATF3* intron 1 and exon 2. The PCR products were cloned into pCR2.1 Topo vector (Invitrogen, #K450002) and the sequence analyzed by Sanger sequencing to verify the KO.

### ZIKV Infection

Twenty-four hours prior infection, cells were seeded in a 100mm tissue culture dish at 1 x10^6^ cells/dish for WT cells and 1.2 x 10^6^ cells/dish for ATF3 KO cells. At this cell density, the cells were near 80% confluent on day of infection. Control cells were trypsinized and counted to determine the volume of virus required for a moi of 1 or 10 plaque forming units (PFU)/cell. An aliquot of viral stock was then thawed at RT, and an appropriate volume of the viral stock was diluted in PBS (Gibco, #14190250) to a final volume of 1 ml and added to cells. For mock-infected plates, 1 ml of PBS was added. Cells were incubated at 37°C for 1 hour, rocking every 15 minutes. An hour later, 9 ml of media was added per plate and returned to the incubator for 24 hours.

### siRNA, shRNA, and Poly I:C Transfections

Single stranded oligos synthesized by Integrated DNA Technologies (IDT) were used for transient transfections. Sense (5’-CGUACGCGGAAUACUUCGAUU-3’) and anti-sense (5’-UCGAAGUAUUCCGCGUACGUU-3’) oligos targeting the control *Gaussia* luciferase GL2 gene (104), were prepared by incubating in annealing buffer (150 mM Hepes [pH 7.4], 500 mM potassium acetate, and 10 mM magnesium acetate) for 1 minute at 90°C followed by a 1-hour incubation at 37°C. The duplex had a final concentration of 20 µM. Prior to transfection, 4 x 10^5^ A549 cells were seeded in 6-well plates for 24 hours. The cells were then transfected with 50 nM control and ATF4 SilencerSelect siRNA (ThermoFisher Scientific, #s1702) using Lipofectamine RNAi Max transfection reagent (Invitrogen, #13778100) based on the manufacturer’s protocol.

To generate A549 cells stably expressing shRNAs, the following the lentivirus approach was performed. HEK 293FT cells were transfected with 1μg of TRC-pLKO.1-Puro plasmid containing either non-targeting shRNA (5’-CAACAAGATGAAGAGCACCAA-3’) or ATF4-targeted shRNA (5’-GCCTAGGTCTCTTAGATGATT-3’) (Sigma-Aldrich), together with 1 μg mixture of packaging plasmids (pMD2.G and psPAX2) prepared in JetOptimus reagent and buffer (Polyplus, #101000025) as per the manufacturer’s instructions. After 24 and 48 hours of transfection, media containing lentivirus was harvested, pooled together, and filtered through a 0.45 µm filter. Pre-seeded A549 cells were subsequently transduced with the lentivirus in the presence of 6 μg/ml of polybrene (Sigma-Aldrich, TR1003). After 24 hours, the lentivirus-containing media was removed, replaced with fresh media and cells were incubated at 37°C for 24 hours. Following incubation, the transduced cells were transferred into new tissue culture dishes and puromycin (1 μg/ml) (InvivoGen, #ant-pr-1) selection was carried out for 4 days. Finally, we screened the transfected and transduced cells by western blot and RT-qPCR to assess the efficiency of knockdown.

A549 WT and ATF3 KO cells were transfected with 1 μg/ml Poly I:C (Sigma-Aldrich, #P1530) for 6 hours at 37°C using Lipofectamine 3000 transfection reagent (Invitrogen, #L3000015) (105). Cellular RNA and proteins were harvested after transfection for further analysis.

### Chemical Treatments

Tunicamycin (Sigma-Aldrich; #T7765) was dissolved in DMSO (Sigma-Aldrich, #34869) at a stock concentration of 2 mM. ER stress was induced by treating cells with 2 μM tunicamycin for 6 hours at 37°C. GSK2606414 (PERK inhibitor; Sigma-Aldrich, #516535) was dissolved in DMSO (Sigma-Aldrich, #34869) to achieve a 30 μM stock concentration. Cells that were mock and ZIKV infected were co-treated with PERK inhibitor at a final concentration of 30 nM for 24 hours at 37°C. ISR Inhibitor (ISRIB; Sigma-Aldrich, #SML0842) (54–56), was reconstituted at 5 mM stock concentration in DMSO (Sigma-Aldrich, #34869) and used at 500 nM on cells for 24 hours at 37°C. Ruxolitinib, a selective inhibitor of JAK 1/2 was reconstituted in DMSO to a stock concentration of 10 mM. Mock and ZIKV-infected cells were simultaneously treated with Ruxolitinib (Selleckchem, #S1378) at 30 nM for the duration of infection. Cells were stimulated with 10 ng/ml IFN-β (R&D Systems, #8499-IF-010) diluted in sterile water for 24 hours at 37°C.

### Harvest of Chemically Treated and ZIKV-Infected Cells

Mock- and virus-infected and chemically treated cells were harvested as follows; first media was aspirated from the cell culture dishes. Cells were gently washed twice with 4 ml cold PBS (Gibco, #14190250) and aspirated. A volume of 1 ml cold PBS (Gibco, #14190250) was then added to the plates, cells were scraped off the plate using a cell lifter and the cell suspension was thoroughly mixed. Equal volumes of 500 μl were aliquoted into two separate tubes. Cell suspensions were centrifuged at 14,000 rpm for 30 seconds to pellet the cells. The supernatant was aspirated off and cells in one tube were prepared for protein analysis while the other tube was prepared for RNA analysis.

### Cell Viability Assay

A549 cells in a 96-well plate were seeded at 4x10^3^ cells/well in 100 μl media and incubated at 37°C 2 days prior to cell viability measurements. Next, cells were treated with the pharmacological inhibitor (GSK2606414 or ISRIB) in 100 μl of media and incubated at 37°C. After 24 hours, plates were removed from incubator and allowed to equilibrate to room temperature for 30 minutes. A volume of 100 μl of CellTiter-Glo 2.0 reagent (Promega, #G9241) was then added to each well and mixed on an orbital shaker for 2 minutes to lyse the cells. The plate was incubated in the dark for 10 minutes to stabilize the signal and the luminescence was read using a Promega GloMax 96 Microplate Luminometer. Cell viability data were obtained from three biological replicates.

### Western Blot Analysis

Cells were lysed with RIPA buffer (100 mM Tris-HCl pH 7.4, 0.1% sodium dodecyl sulphate (SDS), 1% Triton X-100, 1% deoxycholic acid, 150 mM NaCl) containing protease and phosphatase inhibitors (EDTA-free; ThermoScientific, #A32961) and incubated on ice for 20 minutes. The lysates were centrifuged at 14,000 rpm for 20 minutes at 4°C and the clarified supernatant collected. Protein concentrations were quantified using the DC protein assay kit (Bio-Rad, #5000111EDU). Twenty-five micrograms (25 μg) of proteins were separated in 8%, 10% or 12% SDS-polyacrylamide (PAGE) gel at 100 V for 2 hours. Proteins from gels were transferred on to polyvinylidene difluoride membrane (Millipore, #IPVH00010) at 30 V overnight, 100 V for 1 hour or 70 V for 45 minutes at 4°C, respectively. The blots were activated in absolute methanol (Phamco-Aaper, #339000000) and stained with PonceauS (Sigma-Aldrich, #P7170) to determine transfer efficiency. Next, blots were washed in PBS buffer (Gibco, #14190250) with 0.1% Tween (Sigma-Aldrich, #P7949) and blocked in 5% milk or 5% BSA (Sigma-Aldrich, #A9647) in PBS-T for 1 hour at room temperature. The blots were incubated with primary antibodies diluted in blocking buffer for 1 or 2 hours at room temperature or overnight at 4°C. This was followed with three 10-minute PBS-T washes after which the blots were incubated in secondary antibodies diluted with blocking buffer for 1 hour at room temperature. The blots were washed three times in PBS-T and the proteins were visualized using Clarity Western ECL blotting substrate (Bio-Rad, #1705061) or SuperSignal West Femto (ThermoScientific, #34094). The following primary antibodies were used: rabbit anti-ZIKV NS1 (GeneTex, GTX133307; 1:10,000), mouse anti-GAPDH (ProteinTech, #60004-1-lg; 1:10,000), rabbit anti-ATF3 (Abcam, #AB207434; 1:1,000), rabbit anti-ATF4 (D4B8) (Cell Signaling, #11815; 1:1,000), rabbit anti-PERK (D11A8) (Cell Signaling, #5683; 1:1,000), rabbit anti-eIF2⍺ (D7D3)(Cell Signaling, #5324; 1:1,000), rabbit anti-p-eIF2⍺ (D9G8) (Cell Signaling, #3398; 1:1,000), rabbit anti-STAT1 (D1K9Y) (Cell Signaling, #14994; 1:1,000), rabbit anti-phospho-STAT1 (D4A7) (Cell Signaling, #7649; 1:1,000), rabbit anti-STAT2 (D9J7L) (Cell Signaling, #72604; 1:1,000), rabbit anti-phospho-STAT2 (D3P2P) (Cell Signaling, #88410, 1:1,000), rabbit anti-IFIT1 (D2X9Z) (Cell Signaling, #14769; 1:1,000), rabbit anti-MX1(D3W7I) (Cell Signaling, #37849, 1:1,000), rabbit anti-fibrillarin (Abcam, #Ab166630, 1:6000), mouse α-tubulin (Proteintech, #,66031-1-Ig, 1:5,000). Donkey anti-rabbit-IgG (Invitrogen, #31458) and donkey anti-mouse-IgG-HRP (Santa Cruz Biotech, #sc-2314) were used as secondary antibodies at a 1:10,000 dilution. In Figure 6K and Figure 6L, we show the same PVDF membrane that was probed for STAT1, IFIT1 and GAPDH. This blot is denoted by #. The images have been separated into the two figures panels.

### Plaque Assays

Vero cells were seeded in 6-well plates at a density of 7x10^5^/well and incubated at 37°C with 5% CO_2_ overnight. The following day, ten-fold serial dilutions from 10^-1^ to 10^-6^ of media from infections were prepared in 1 x PBS (Gibco, #14190250). The media on Vero cells seeded the previous day was aspirated, 150 μl of 1 x PBS was added to the mock well, and 150 μl of each virus dilution was added to the remaining wells. The cells were incubated at 37°C with 5% CO_2_ for 1 hour, with gentle rocking every 15 minutes. After incubation, the PBS or virus dilution in PBS was aspirated and 3 ml of overlay consisting of 1:1 2 x DMEM (DMEM high glucose, no sodium bicarbonate buffer powder [Gibco # 12-100-046] in 500 mL of RNase-free water, 84 mM of sodium bicarbonate, 10% FBS and 2% penicillin and streptomycin, at pH 7.4) and 1.2% avicel (FMC, #CL-611) was added to each well and the plates were incubated at 37°C with 5% CO_2_. Five days post-infection, the overlay was aspirated, cells were fixed with 1 ml of 7.4% formaldehyde (Fischer Scientific, #F79-500) for 10 minutes at room temperature, rinsed with water and plaques were visualized using 1% crystal violet (ThermoScientific, #R40052) in 20% methanol. Viral titers were determined from duplicate viral dilutions and three biological replicates.

### RT-qPCR Analysis

Total RNA was isolated from cells using TRIzol reagent (Invitrogen, #15596026) and the RNA Clean and Concentrator kit (Zymo Research, #R1018). The RNA was DNase-treated using the TURBO DNA-free^TM^ kit (Invitrogen, #AM1907) and reverse transcribed using the High-Capacity cDNA Reverse Transcription reagents (Applied Biosystems, #4368813). The resulting cDNA was used for qPCR analysis with iTaq Universal SYBR Green Supermix reagents (Biorad, #1725124) and CFX384 Touch Real-Time PCR system (Biorad). RT-qPCR data shown are from at least three independent experiments, with each sample assayed in three technical replicates. The RT-qPCR primer sequences are shown in Table 1.

**Table 1:**
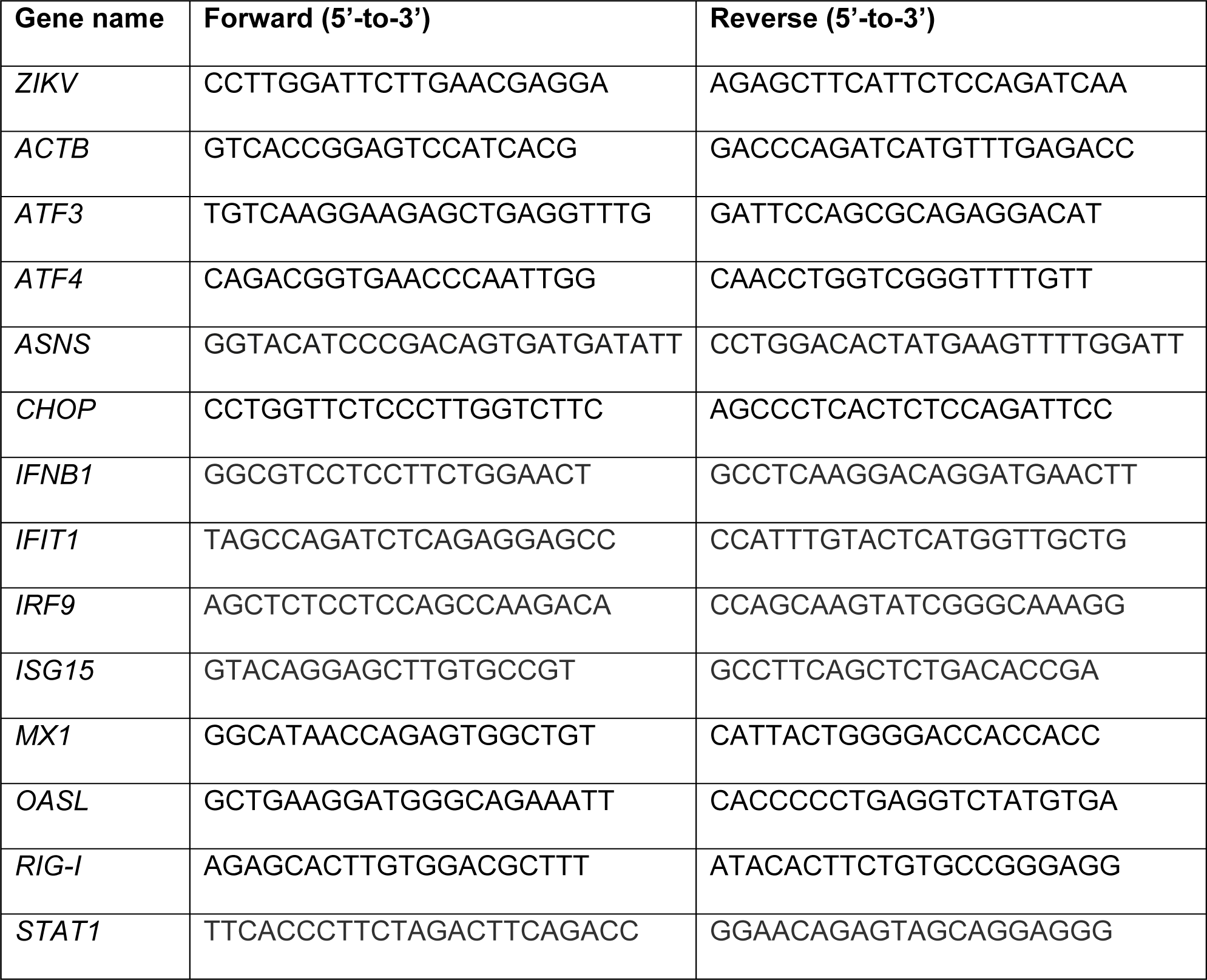
Primers used for RT-qPCR.

### Statistical Analysis

The data shown are from at least three independent experiments. Data were analyzed using Prism 9.4.1 software (GraphPad, La Jolla, CA, USA) to establish statistical significance. We performed two-tailed student T-test for two group comparisons.

### RNA-seq sample processing and analysis

A549 WT and ATF3 KO cell lines were either mock or ZIKV infected at a moi of 10, as described above. At 24 hours post-infection, cells were harvested, and total RNA was isolated using TRIzol reagent (Ambion by Life Technologies) and the RNA Clean and Concentrator kit (Zymo Research, #R1018). Total RNA was DNAse-treated with the TURBO DNAse-free^TM^ reagent (Invitrogen, #AM1907) and RNA quality was assessed via Bioanalyzer 2100 RNA analysis. Only samples with an RNA Integrity Number (RIN) greater than 8.5 were used for subsequent experiments. PolyA-selected, strand-specific RNA-seq libraries were generated and sequenced in paired-end mode (150 x 2) on an Illumina HiSeq 3000 by Genewiz (Azenta Life Sciences). Raw FastQ files and DESeq2 results tables are deposited in Gene Expression Omnibus via accession number GSE233049.

### Differential Gene Expression Analysis

Abundance of transcripts from the *Ensembl* hg38 genome/transcriptome assembly (v.104) was quantified using *kallisto* in quant mode with 100 bootstraps (106). Transcript counts (in TPM, transcripts per million) were imported into the *R* statistical computing environment via *tximport* (107). Differential gene expression between infection and genotype conditions were quantified using DESeq2 (108). Principal component analysis (PCA) was performed on the top 4000 transcripts with the highest expression levels (TPM).

### Gene ontology analysis

Gene Ontology (GO) terms and enrichment statistics were derived from performing GO analysis using Metascape (109). Gene lists were extracted from DESEQ2 (108) results comparing between genotypes and treatment conditions (WT ZIKV vs MOCK, ATF3 KO MOCK vs ZIKV, MOCK ATF3 KO vs WT or ZIKV ATF3 KO vs WT) where changes in expression were significant (padj > 0.05) and substantial (2-fold change [2FC], upregulated or downregulated). A single gene list for every genotype and treatment combination was used as an input for Metascape offline analysis with Reactome and default search parameters. Metascape Gene Ontology terms and associated statistics were used as an input to generate dotplots with top 10 terms for each sample with GSEApy library (110). Heatmaps in Fig. 6F were generated using DESEQ2 normalized counts which were row-wise normalized using z-score.

## Abbreviations

ATF3: Activating transcription factor 3
ATF4: Activating transcription factor 4
BMDMs: Bone marrow-derived macrophages
CHOP: C/EBP homologous protein
DENV: Dengue virus
DMSO: Dimethyl sulfoxide
eIF2⍺: Eukaryotic initiation factor 2-alpha
GCN2: General control non-derepressible-2
HRI: Heme-regulated eIF2⍺ kinase
IFN: Interferon
ISG: Interferon stimulated genes
ISR: Integrated stress response
ISRIB: Integrated stress response inhibitor
JEV: Japanese encephalitis virus
MCMV: murine cytomegalovirus
NS: Nonstructural
PKR: Protein kinase R; double-stranded RNA-dependent protein kinase
PERK: Protein kinase R-like ER kinase
UPR: Unfolded protein response
ZIKV: Zika virus
ZIKV PRVABC59: Zika virus Puerto Rico isolate
ZIKV MR766: Zika virus Ugandan isolate

## Data Availability

RNA-seq data from wild-type and ATF3 knockout A549 human lung adenocarcinoma cells either mock-infected or infected with ZIKV PRVABC59 at a moi of 10 PFU/cell and harvested at 24 hours post-infection has been deposited in Gene Expression Omnibus (GEO) at GSE233049.

## Acknowledgments

This work was supported by grants from National Institutes of Health to CTP (R01GM123050 and R21AI178672) and MAS (R35GM138120). PB was supported by a generous predoctoral fellowship from the American Heart Association (Award ID: 903514). The research in this manuscript is solely the responsibility of the authors and does not necessarily represent the official views of the NIH or AHA. We also gratefully acknowledge Kristen Kaytes, and Drs. Marlene Belfort and John Cleary at UAlbany and The RNA Institute for their thoughtful comments and suggestions on this manuscript.

## Notes

### Competing Interest Statement

The authors have declared no competing interest.

### Summary of Updates

Data that was in the supplement has been incorporated into the main figures. There are now no supplemental data. Figure 7 from the first version of the manuscript has been removed. These data showed the role of ATF3 in autophagy. Three new figures are included in the revised manuscript (Figures 6-8). These new data include transcriptomic analysis (Figure 6), activation of the innate immune response pathway in wild-type and ATF3 knockout cells following poly I:C and interferon-beta treatment (Figure 7), and effects on Zika virus infection in wild-type and ATF3 knockout cells when treated with a JAK inhibitor (Figure 8). The text has been modified to reflect these changes.

